# Multiscale Functional Dysconnectivity Reveals Sex-Dominant Neurosubtypes and Predicts Unique Neurocognitive Pathways in Psychosis

**DOI:** 10.64898/2026.09.04.749545

**Authors:** Ram Ballem, Pablo Andrés-Camazón, Covadonga M. Díaz-Caneja, Juan R Bustillo, Jessica A. Turner, Adrian Preda, Vince D. Calhoun, Jiayu Chen, Armin Iraji

**Affiliations:** Department of Computer Science, Georgia State University, Atlanta, USA; School of Electrical and Computer Engineering, Georgia Institute of Technology, Atlanta, USA; Tri-Institutional Center for Translational Research in Neuroimaging and Data Science (Georgia State University, Georgia Institute of Technology, Emory University), Atlanta, USA; Department of Child and Adolescent Psychiatry, Institute of Psychiatry and Mental Health, Hospital General Universitario Gregorio Marañón, Instituto de Investigación Sanitaria Gregorio Marañón (IiSGM), CIBERSAM, ISCIII, School of Medicine, Universidad Complutense, Madrid, Spain; Department of Psychiatry and Behavioral Sciences, University of New Mexico, Albuquerque, New Mexico; Department of Psychiatry and Behavioral Health, Wexner Medical Center, The Ohio State University, Columbus, OH, USA; Department of Psychiatry and Human Behavior, University of California, Irvine, California, USA

**Author notes:** Corresponding Author: Name: Ram Ballem, Postal Address: 55 Park Place NE, 18^th^ Floor – TReNDS Center Atlanta, GA – 30303, Name: Armin Iraji, Postal Address: 55 Park Place NE, 18^th^ Floor – TReNDS Center, Atlanta, GA – 30303. Contributing equally (Jiayu Chen, Armin Iraji).

## Abstract

Psychotic disorders are severe mental conditions whose biological underpinnings remain elusive. This is in part due to marked heterogeneity, which may have precluded advances in precise understanding and treatment of the disorders. Previous work has tried to elucidate this heterogeneity; nevertheless, sex, as an important biological variable, has not been well accounted for despite established sex differences. Using resting-state fMRI data from the BSNIP consortium (N=1753; 64.3% probands with psychosis include 571 females and 556 males; 35.7% of controls includes 369 females and 257 males), we extracted multiscale functional network connectivity and employed unsupervised learning methods to identify potential subgroups with distinct neurobiological profiles, characterized by an overrepresentation of males or females (termed here as sex-dominant Psychosis Imaging Neurosubtypes, PINs).

Four sex-dominant neurosubtypes emerged, two female-dominant (fPIN-1, fPIN-2) and two male-dominant (mPIN-1, mPIN-2). Each PIN exhibited distinct, replicable dysconnectivity patterns that uniquely contributed to cognitive deficits. Brain-Predicted Cognitive performance derived from these PIN-specific dysconnectivity, prominent among the triple network (default mode and salience), subcortical-basal ganglia, higher cognition-insular temporal, and visual-occipitotemporal subdomains, correlated with measured cognitive performance and showed PIN-specific reductions relative to controls, confirming neurosubtype-congruent associations. Moreover, fPIN-2’s distinct dysconnectivity also predicted positive and negative symptom scores.

Our findings lend support for sex-dominant neurobiological heterogeneity, with sex-dominant PINs presenting unique dysconnectivity patterns that contribute to cognitive and symptom outcomes, even though neurosubtypes appear clinically similar. These results establish sex as a critical biological variable in deconstructing psychosis heterogeneity, which holds promise in revealing more precise biomarkers for disease characterization and guiding personalized treatment strategies.

## 1. INTRODUCTION

Psychosis encompasses a spectrum of severe mental disorders, characterized by delusions and/or hallucinations and disorganized thinking and behavior^1^. Schizophrenia (SZ) is the most common and severe form of chronic psychotic disorders^2^, with a lifetime prevalence of about 1%^3^ and a substantial economic burden of more than $150 billion annually just in the United States^3^. Other chronic psychotic disorders, such as Schizoaffective Disorder (SAD) and Psychotic Bipolar Disorder, present lower prevalence (0.5-0.8%) and show considerable clinical and biological overlap between them and with SZ, supporting the view that they exist on a continuum^4,5^. Unfortunately, treatment outcomes remain poor, with nearly 40% of individuals showing limited response to medication^6,7^. A major barrier to therapeutic progress is pronounced biological heterogeneity underlying psychosis, which likely reflects multiple interacting mechanisms. Importantly, sex represents an important but unexplored dimension of this heterogeneity^8^, highlighting the need for a deeper understanding of underlying biology to facilitate more effective and precise treatment for psychotic disorders.

Psychotic disorders are biologically heterogeneous conditions, and this heterogeneity is likely driven by multiple underlying mechanisms or their various combinations^9,10^. Traditional case-control approaches based on Diagnostic and Statistical Manual of Mental Disorders (DSM) diagnoses may not fully capture this underlying biological heterogeneity, leaving it unclear whether the disorder represents a single illness or different illnesses that lead to similar clinical presentations. This uncertainty continues to limit progress towards biomarkers and precision interventions^11^. To address this, neuroimaging studies, leveraging machine learning, have increasingly sought to uncover biological subgroups (hereafter referred to as imaging neurosubtypes), with replicable subtypes linked to cognitive, symptom, and illness course^12–24^. These approaches vary in methodology, drawing on features either associated with outcomes or those that differentiate patients from controls. In most subtyping studies, sex has been statistically balanced in sample design or controlled as a covariate in multivariate models, rather than explicitly investigated as another biologically meaningful dimension. Studying its role in psychosis heterogeneity is essential, as growing evidence indicates that sex exerts a fundamental influence on the neurobiology of psychosis.

Sex differences in psychosis have been recognized across prevalence, age of onset, symptomatology, cognitive impairment, and brain function^25–28^, though findings remain variable and sometimes inconsistent. Psychotic disorders^29,30^ are generally considered to be more prevalent in males, although some studies report comparable rates^31^, with earlier onset and more severe negative symptoms. Females more often show later onset, more affective and positive symptoms, and initially better clinical profiles that tend to converge with those of males over time^32,33^. Cognitive outcomes are equally heterogeneous, with reports ranging from better performance in females to advantages in males, to no sex-based differences at all^26^. Neuroimaging studies also suggest sex-specific neurobiology. Resting-state functional magnetic resonance imaging (rsfMRI) has shown altered frontal blood oxygen level-dependent (BOLD) activity in males and females with SZ, with sex-dependent differences in neuronal activity and metabolic demands that may contribute to distinct clinical manifestations^34^. Other studies report greater subcortical and posterior region activity in females, some of which correlate with depressive symptoms^35^. Sex-specific alterations in sensorimotor and temporal brain regions have been linked to positive and negative syndrome scale (PANSS) scores^25^. Additionally, functional network connectivity (FNC) differences between the triple network domain, including central executive, salience, and default mode networks, have been linked to genetic risk of SZ in both males and females^36^, and symptom severity in males but not in females^25^. Collectively, these findings highlight both the complexity of psychosis and the limitation of not accounting for sex when deconstructing heterogeneity in psychosis.

Different biological reasons could explain why certain neurosubtypes may be much more prevalent in one sex than the other. Males and females with psychosis develop under distinct genetic, biological, and environmental backgrounds^31^. In females, estrogen and other sex hormones exert neuroprotective efforts^26,28^, potentially contributing to later onset and a more favorable illness course. Genetic and epigenetic factors, including SNPs and DNA methylation, have also been suggested to contribute to sex-specific onset patterns^37^. Risk and protective factors for psychiatric disorders are associated with structural and functional brain patterns, and these factors often differ between males and females^38–41^. Environmental exposures, including adolescent cannabis use^42,43^ and childhood trauma^33,44^, that have been linked with the development of psychosis, also follow sex-specific patterns in boys and girls that potentially shape brain development in a sex-specific manner^45^.

Contradictory findings on sex differences in psychosis may be explained by the presence of sex-dominant heterogeneity, defined here as patterns of neurobiological variation that are enriched in one sex but not exclusive to it, which lead to differential brain function between men and women. This heterogeneity has been largely overlooked in previous studies and may blur the true biological effects. Accounting for this heterogeneity and identifying neurosubtypes that capture distinct sex-dominant neurobiological signatures could enable stratification beyond DSM diagnoses and improve our understanding of how sex shapes the neurobiological architecture and clinical manifestation of psychosis. We hypothesize that there exist sex-dominant neurosubtypes with male or female overrepresentation, but not exclusively confined to them, which reflects the stratification along the spectrum of male-dominant to female-dominant brain alterations with shared characteristics^46,47^.

In this study, we test the hypothesis that sex-related neurobiological heterogeneity in psychosis manifests as distinct but sex-dominant neurosubtypes, rather than sex-exclusive ones. Using rsfMRI, we investigate the presence of sex-dominant psychosis imaging neurosubtypes (PINs) exhibiting distinct functional connectivity patterns that may differentially relate to symptomatology and cognitive performance. To this end, we analyze a large cohort including probands with SZ, SAD, and Bipolar I Disorder with psychotic features (BP) and healthy controls. Leveraging the standardized Neuromark 2.2 template, which incorporates 105 multiscale intrinsic connectivity networks (ICNs) derived from a large population^48^, we extracted subject-specific multiscale functional network connectivity (msFNC). Unsupervised models were designed to identify connectivity patterns that differ between males and females, while allowing individuals of either sex to express these patterns. This process in an ensemble framework provides a robust foundation for subgrouping analyses^13,49^ and facilitates the identification of clinically relevant, sex-dominant PINs. Dimensionality reduction followed by ensemble learning was subsequently applied to delineate these robust PINs and characterize their cognitive and symptom associations.

## 2. RESULTS

### 2.1. Sex-dominant PINs Identification and Replication

We analyzed rsfMRI from the Bipolar-Schizophrenia Network on Intermediate Phenotypes (B-SNIP; N = 1753; probands with psychosis = 1127 and controls = 626; see Methods 7.1)^51^. Using the robust Neuromark pipeline with the standardized NeuroMark 2.2 multiscale template^48,50^, we extracted 105 canonical and replicable msICNs (see Methods 7.2). For each participant, we quantified subject-level msFNC by computing Pearson correlations between msICN time courses (Figure 1, left panel), yielding a 105×105 connectivity matrix. The dataset was divided into a discovery set (∼80% of the sample) and a replication set (∼20% of the sample), with demographic, symptom, and cognitive characteristics summarized in Table 1.

**Figure 1.**
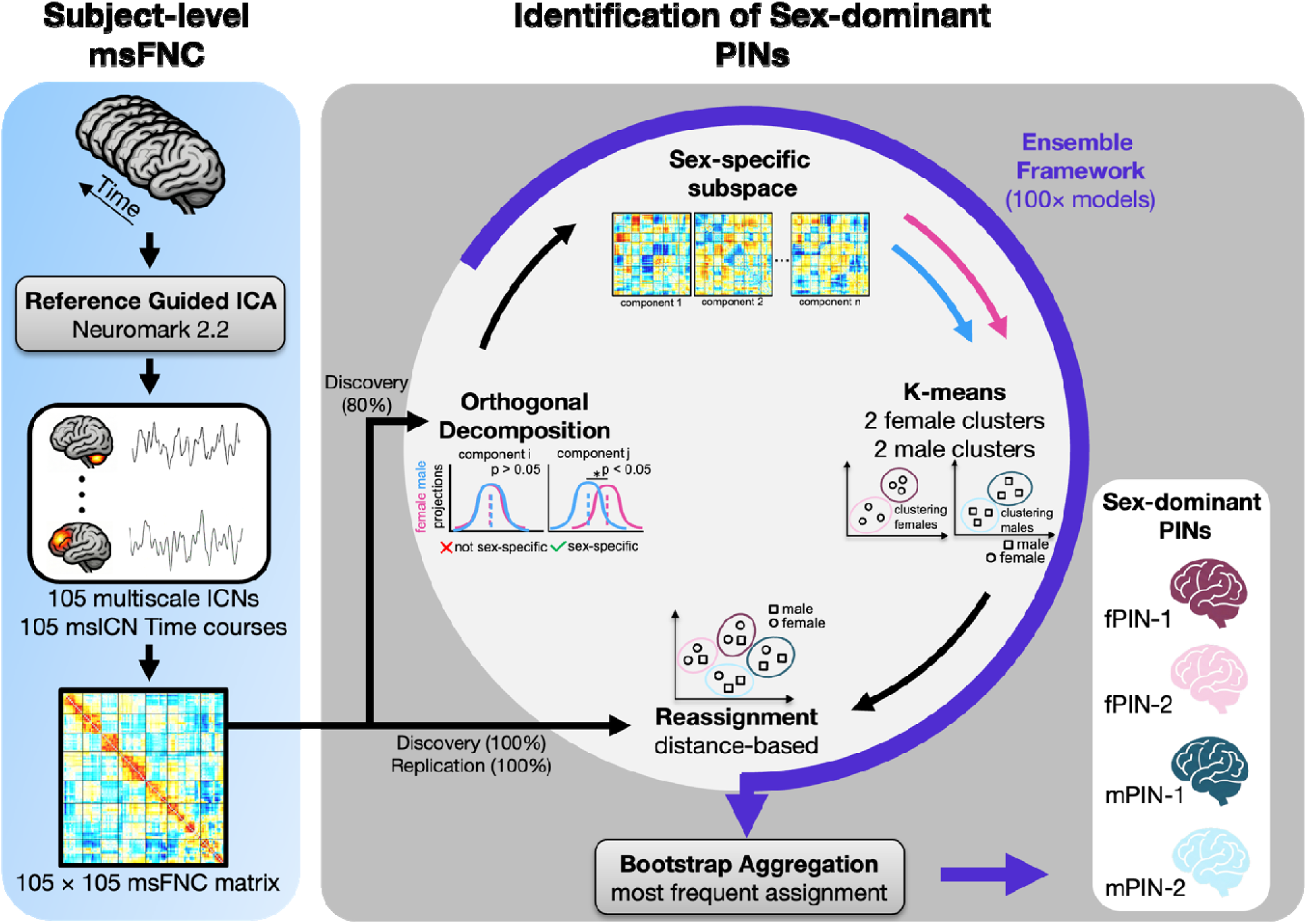
Pipeline for multiscale Functional Network Connectivity (msFNC) estimation and sex-dominant Psychosis Imaging Neurosubtypes (PINs) identification. Left Panel: Resting-state functional magnetic resonance imaging (rsfMRI) data for a single subject were processed through the Neuromark 2.2 pipeline^48,50^, yielding 105 multiscale intrinsic connectivity networks (ICNs) and their time courses. These were used to compute a whole-brain 105 × 105 msFNC matrix (see Methods 7.2). Right Panel: Principal component analysis (PCA) was applied to the msFNC matrices of psychosis probands (PSY), and Principal components (PCs) showing significant sex differences were identified to define a sex-specific subspace. Projections onto this subspace were clustered separately for males and females, followed by reassignment across all subjects based on minimum distance criteria. An ensemble bootstrap aggregation framework (100 models) was used to enhance cluster stability, resulting in two female-dominant (fPIN-1, fPIN-2) and two male-dominant (mPIN-1, mPIN-2) PINs. This framework enables the identification of stable and reliable sex-dominant connectivity-based neurosubtypes in psychosis.

**Table 1.** Demographic, symptom, and cognitive characteristics.

|  | Discovery set (Train set) |  | Replication set (Test set) |  |
| --- | --- | --- | --- | --- |
|  | Controls | Probands | Controls | Probands |
| **Demographics mean±sd** |  |  |  |  |
| N (%) | 500 (35.95%) | 891 (64.1%) | 126 (34.8%) | 236 (65.2%) |
| Male N (%) | 206 (41.2%) | 454 (50.95%) | 51 (40.48%) | 102 (43.22%) |
| Female N (%) | 294 (58.8%) | 437 (49.05%) | 75 (59.52%) | 134 (56.78%) |
| Age | 35.92±12.20 | 36.58±12.22 | 35.55±12.94 | 37.33±11.84 |
| **Clinical Symptoms mean±sd** |  |  |  |  |
| Young Mania Rating Scale (YMRS) | NA | 7.99±7.45 | NA | 8.41±6.68 |
| Montgomery and Asberg Depression Rating Scale (MADRS) | NA | 10.75±9.52 | NA | 12.46±9.82 |
| Positive and Negative Syndrome Scale (PANSS) positive | NA | 15.67±6.06 | NA | 15.63±5.77 |
| PANSS negative | NA | 14.64±6.38 | NA | 14.77±6.38 |
| PANSS general | NA | 30.68±5.98 | NA | 31.19±6.38 |
| PANSS total | NA | 60.98±18.64 | NA | 61.57±19.11 |
| **Cognitive assessment mean±sd** |  |  |  |  |
| Brief Assessment of Cognition in Schizophrenia - composite score (BACS-COMP) | -0.26±1.21 | -1.49±1.39 | -0.17±1.18 | -1.45±1.41 |
| **Race N (%)** |  |  |  |  |
| African American | 167 (33.4%) | 339 (38.0%) | 33 (26.2%) | 91 (38.6%) |
| American Indian | 2 (0.4%) | 3 (0.3%) | 0 (0%) | 1 (0.4%) |
| Asian | 35 (7%) | 24 (2.7%) | 8 (6.3%) | 9 (3.8%) |
| Caucasian | 264 (52.8%) | 451 (50.6%) | 75 (59.5%) | 113 (47.9%) |
| More than one race | 16 (3.2%) | 48 (5.4%) | 6 (4.8%) | 15 (6.4%) |
| Other | 14 (2.8%) | 26 (2.9%) | 4 (3.2%) | 7 (3%) |
| Pacific Islander | 2 (0.4%) | 0 (0%) | 0 (0%) | 0 (0%) |
| **Site N (%)** |  |  |  |  |
| Baltimore | 42 (8.4%) | 102 (11.4%) | 9 (7.1%) | 18 (7.6%) |
| Boston | 26 (5.2%) | 65 (7.3%) | 8 (6.3%) | 15 (6.4%) |
| Chicago | 113 (22.6%) | 225 (25.3%) | 32 (25.4%) | 58 (24.6%) |
| Dallas | 96 (19.2%) | 139 (15.6%) | 18 (14.3%) | 44 (18.6%) |
| Detroit | 15 (3%) | 35 (3.9%) | 7 (5.6%) | 15 (6.4%) |
| Georgia | 81 (16.2%) | 86 (9.7%) | 22 (17.5%) | 22 (9.3%) |
| Hartford | 127 (25.4%) | 239 (26.9%) | 30 (23.8%) | 64 (27.1%) |

To mitigate potential age-related confounding, we statistically adjusted for age effects before dimensionality reduction, ensuring that the identified subtypes were not driven by age-related variations in msFNC. We applied principal component analysis (PCA) to the discovery set msFNC data from probands (Figure 1, right panel) and retained the top principal components (PCs) explaining 99% of the variance. Two-sample t-tests on this retained PCA projection identified PCs with significant sex effects, which were then used for clustering. Clustering analysis was performed separately on female and male projections and consistently identified two optimal clusters for each sex group. Next, we relaxed the sex constraint and assigned individuals to one of the four clusters based on minimum distance. While sex-specific clustering enhances the ability to identify distinct subgroups within each sex, the relaxed assignment allows individuals from different sexes to be assigned to the most neurobiologically similar cluster. These same clusters were also used to assign individuals in the hold-out replication set based on minimum distance assignment.

To ensure stability and reproducibility, we implemented an ensemble bagging framework (Figure 1, right panel), generating 100 bootstrap models trained on a subsample of the discovery set. Each model produced four cluster centroids (Figure 2.A shows all centroids along the top two PCs with significant sex effects) and relaxed label assignments for subjects. The final label for each subject in both discovery and replication sets was determined by the most frequent cluster assignment across models (see Methods 7.3). This process yielded four representative sex-dominant PINs: female PIN-1 (fPIN-1), female PIN-2 (fPIN-2), male PIN-1 (mPIN-1), and male PIN-2 (mPIN-2). To assess robustness to multisite effects, msFNC was harmonized using neuroCombat and reassigned to the ensemble models. Harmonized labels showed significant concordance with the original assignments (p < 0.001), where concordance was estimated using the Hungarian algorithm and was compared against a non-parametric permutation-based null distribution (see Methods 7.3.3). This result suggests that the identified PINs are not primarily driven by site effects. These ensemble models not only stratified probands in both discovery and replication sets but also provide a transferable framework for assigning sex-dominant PIN membership in other independent datasets.

**Figure 2.**
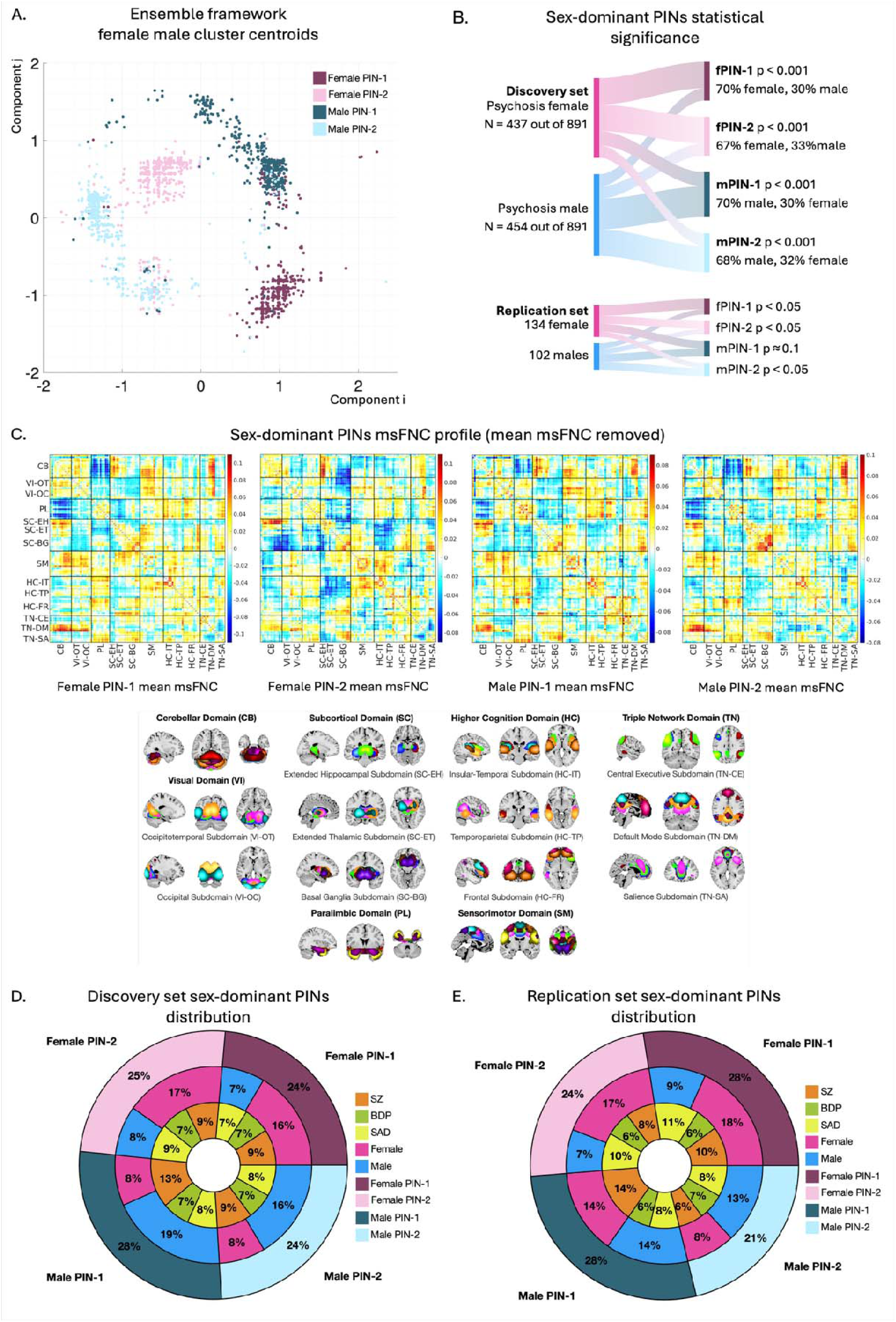
Sex-dominant Psychosis Imaging Neurosubtypes (PINs): msFNC profile, Replication, and Group Distributions. (A) Ensemble framework (100 models) showing female and male cluster centroids along the top two sex-specific components (illustrative subspace, higher dimension in reality). (B) Plots of discovery and replication set assignments, showing sex overrepresentation within sex-dominant PINs, with significance assessed via null distribution and hypothesis testing (p-values shown). (C) msFNC profile (mean msFNC of controls and probands) for fPIN-1, fPIN-2, mPIN-1, and mPIN-2, with corresponding msICNs from B-SNIP participants. msFNC connectivity domains include Cerebellar (CB); Visual (VI), comprising Occipitotemporal (OT) and Occipital (OC) subdomains; Paralimbic (PL); Subcortical (SC), comprising Extended Hippocampal (EH), Extended Thalamic (ET), and Basal Ganglia (BG) subdomains; Sensorimotor (SM); Higher Cognition (HC), comprising Insular Temporal (IT), Temporoparietal (TP), and Frontal (FR) subdomains; Triple Network (TN), comprising Central Executive (CE), Default Mode (DM), and Salience (SA) subdomains. (D, E) Pie charts showing the distribution of sex-dominant PINs in the discovery (D) and replication (E) sets. The outer ring shows PIN distributions, the middle ring shows sex distribution within each PIN, and the inner ring shows DSM diagnoses distribution. These results demonstrate robust replication and clear sex-dominant clustering, supporting the stability and biological relevance of the identified PINs.

To evaluate whether the identified PINs exhibited sex-dominant patterns, we performed non-parametric permutation tests against a null distribution (see Methods 7.3.4). The identified PINs were strongly sex-dominant, with a clear overrepresentation of one sex within each neurosubtype. In the replication set, non-parametric assignment tests (Figure 2.B) validated significant sex overrepresentation in three PINs (fPIN-1: p < 0.05, fPIN-2: p < 0.05, mPIN-2: p < 0.05). mPIN-1 showed a trend toward sex dominance that did not reach the significance threshold (p = 0.14), potentially due to the limited replication sample size. Using these validated labels, we arrived at each PINs msFNC profile by averaging connectivity of neurosubtype groups and removing the mean connectivity of all participants (Figure 2.C).

To determine whether the identified PINs reflect a subtype structure related to psychosis, rather than one that would emerge from applying the same clustering procedure to any general population, we applied the same ensemble framework (Figure 1, right panel; See Methods 7.3) identifying control-derived subtypes. Controls in the replication set were then assigned to these control-derived subtypes, and none showed significant sex overrepresentation (p = 0.15–0.85). This indicates that the strongly sex-dominant subtype structure identified in psychosis does not replicate when the same procedure is applied to controls, supporting that this subtyping result is specific to psychosis rather than a generic outcome of the clustering approach. Distributions of sex-dominant PIN membership, sex, and DSM diagnoses relations are shown in Figures 2.D and 2.E. Additionally, these PIN memberships were also independent of DSM diagnoses (Replication set ^2^ = 7.9, p = 0.24). Collectively, these findings suggest the presence of replicable sex-dominant PINs in psychosis that are not driven by diagnostic categories.

### 2.2. Neurobiological Profile of Sex-dominant PINs

We next examined differences in whole-brain msFNC between each sex-dominant PIN and the control group. Significant replicable dysconnectivity patterns were observed for all four sex-dominant PINs in discovery and replication sets (Figure 3; detailed results in Supplementary Section 4) with adjustment for confounds (race, site, and mean framewise displacement (meanFD); See Methods 7.4). Beyond case-control differences, comparisons between sex-dominant PINs revealed statistically significant msFNC dysconnectivity (Supplementary 4), demonstrating that each PIN has a distinct neurobiological profile. Overall, fPIN-1, fPIN-2, mPIN-1, and mPIN-2 exhibited 460, 511, 748, and 272 replicated dysconnected features, respectively, out of 5460 tested connections (Figure 3, left panels).

**Figure 3.**
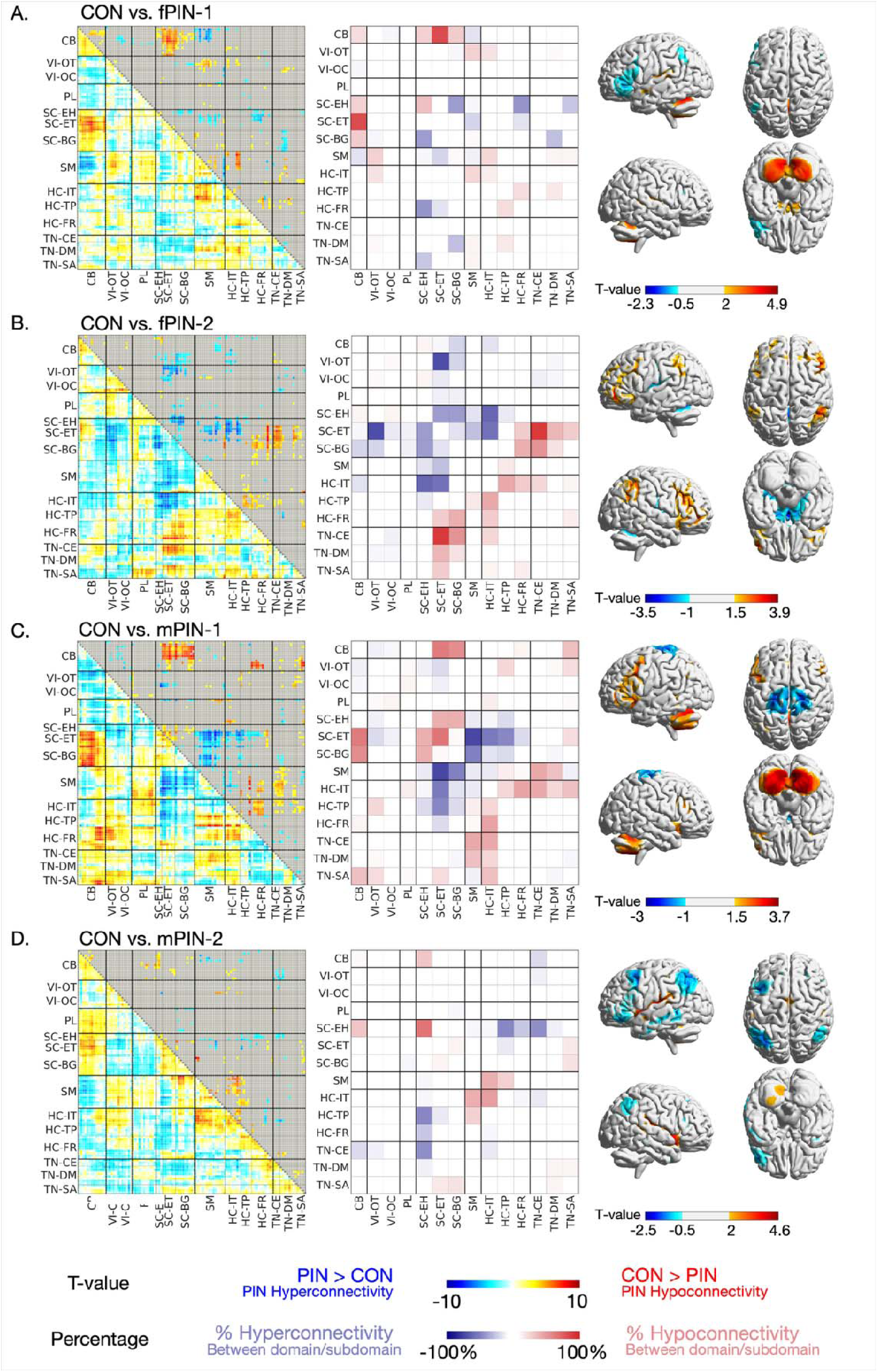
Sex-dominant Psychosis Imaging Neurosubtypes (PINs) multiscale Functional Network Connectivity (msFNC) dysconnectivity. (A) Controls (CON) vs. fPIN-1 msFNC dysconnectivity (B) CON vs. fPIN-2 msFNC dysconnectivity. (C) CON vs. mPIN-1 msFNC dysconnectivity. (D) CON vs. mPIN-2 msFNC dysconnectivity. msFNC dysconnectivity patterns (left panel) were identified for CON vs. sex-dominant PINs, across msFNC. Lower triangles display t-values of group differences, where each cell reflects a group difference in connectivity between two multiscale intrinsic connectivity networks (msICNs). The upper triangle shows statistically significant differences using -log(q-value)*sign(t-value), with non-significant ones grayed out. The center panel heatmaps show dominant significant connectivity differences expressed as percentages across domain-domain/subdomain totals. Surface maps (right panel) of full brain view showing spatial distribution of PINs msFNC dysconnectivity. msFNC connectivity domains include Cerebellar (CB); Visual (VI), comprising Occipitotemporal (OT) and Occipital (OC) subdomains; Paralimbic (PL); Subcortical (SC), comprising Extended Hippocampal (EH), Extended Thalamic (ET), and Basal Ganglia (BG) subdomains; Sensorimotor (SM); Higher Cognition (HC), comprising Insular Temporal (IT), Temporoparietal (TP), and Frontal (FR) subdomains; Triple Network (TN), comprising Central Executive (CE), Default Mode (DM), and Salience (SA) subdomains. These findings reveal that each sex-dominant PIN exhibits a distinct large-scale dysconnectivity pattern relative to controls, highlighting biologically meaningful heterogeneity in psychosis.

fPIN-1 exhibited hyperconnectivity within the subcortical domain and hypoconnectivity between the entire cerebellar domain and subcortical-extended thalamic subdomain (Figure 3.A middle panel). The anatomical regions involved in the dysconnectivity patterns are shown in Figure 3.A right panel.

fPIN-2 showed hyperconnectivity between subcortical-extended thalamic and visual-occipitotemporal, higher cognition-insular temporal subdomains and primarily hypoconnectivity between triple network (central executive, default mode and salience subdomains) and subcortical-extended thalamic subdomains, sensorimotor and higher cognition domains (Figure 3.B middle panel). Surface representations demonstrate visual occipitotemporal and higher cognition networks involved in hyperconnectivity (Figure 3.B right panel).

mPIN-1 displayed major hyperconnectivity between subcortical-extended thalamic subdomain and sensorimotor, higher cognition domains and widespread hypoconnectivity between cerebellar and subcortical domains, higher cognition-frontal and triple network-salience subdomains (Figure 3.C middle panel). Figure 3.C right panel highlights the cerebellar and sensorimotor networks involved in mPIN-1 dysconnectivity.

mPIN-2, while presenting comparatively less dysconnected differences, showed hyperconnectivity primarily between the subcortical-extended hippocampal subdomain and higher cognition, triple network domains (central executive, default mode and salience subdomains), and predominant hypoconnectivity within higher cognition-insular temporal and subcortical extended hippocampal subdomain (Figure 3.D middle panel). These effects are localized to higher cognition, triple network, and cerebellar regions (Figure 3.D right panel).

#### 2.2.1. Sex-dominant PINs distinct dysconnectivity profile

To identify PIN-specific differences, we isolated distinct msFNC dysconnectivity features, defined as connectivity differences unique to a given sex-dominant PIN relative to all other PINs. Uniqueness was defined over statistically significant differences that were replicated across both discovery and replication sets, and a distinct dysconnection for a given PIN. This analysis yielded 225, 341, 488, and 147 for fPIN-1, fPIN-2, mPIN-1, and mPIN-2, respectively (Figure 4, left panels). Collectively, these PIN-specific features define four connectivity disrupted subsystems, each representing a unique profile of network alterations characterizing a particular sex-dominant PIN.

**Figure 4.**
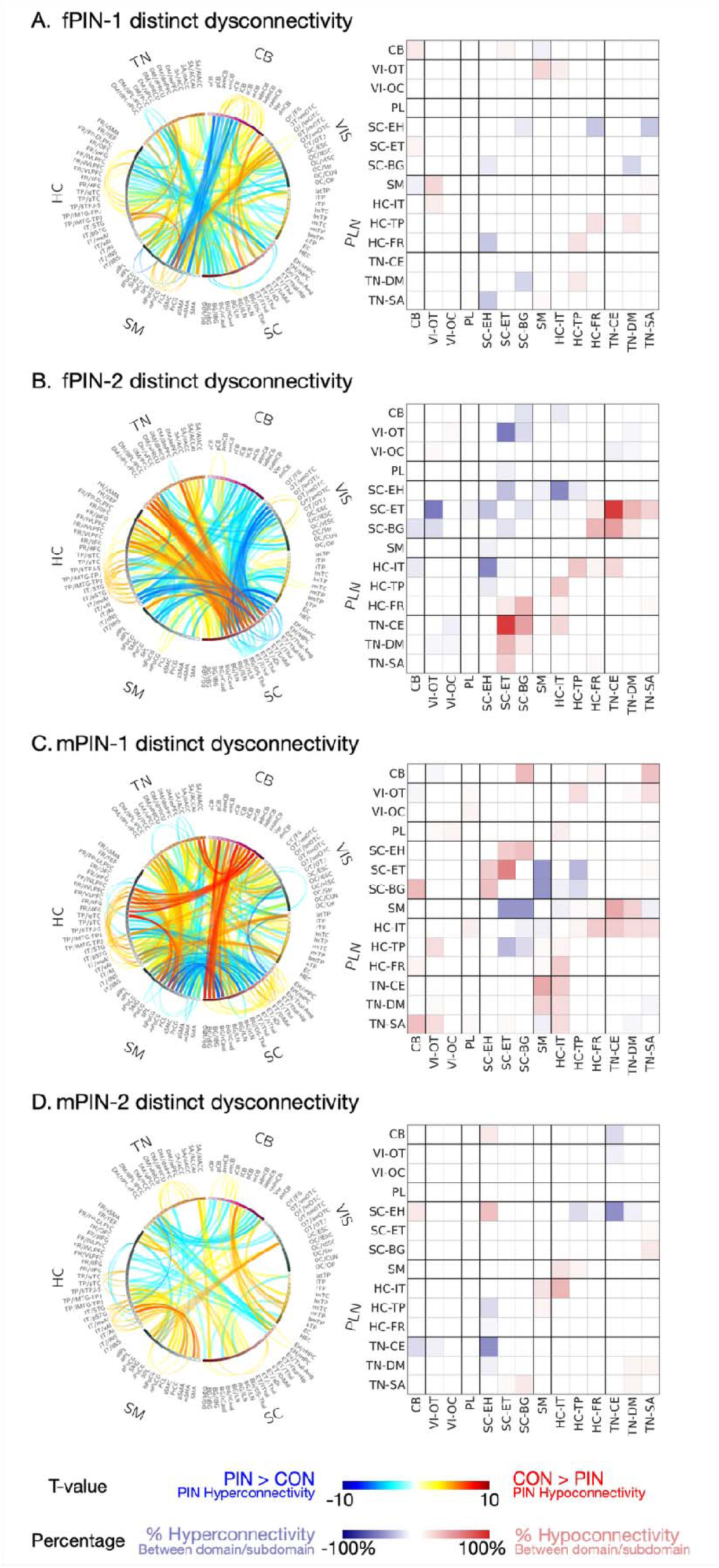
Distinct Dysconnectivity Profile of Sex-dominant Psychosis Imaging Neurosubtypes (PINs). (A) Controls (CON) vs. fPIN-1 distinct (unique) dysconnectivity (B) CON vs. fPIN-2 distinct dysconnectivity. (C) CON vs. mPIN-1 distinct dysconnectivity. (D) CON vs. mPIN-2 distinct dysconnectivity. Distinct (unique) dysconnectivity msFNC features shown as connectograms (left panel) were identified for CON vs. sex-dominant PINs, across msFNC. Connectograms illustrate t-values of group differences, where each link reflects a group difference in connectivity between two multiscale intrinsic connectivity networks (msICNs). Only statistically significant distinct dysconnectivity in discovery, replicated in the hold-out set, is shown. Outward links represent differences in within-domain connectivity, while inward links indicate differences between domains, with link intensity reflecting the t-values. The right panel heatmaps show dominant significant connectivity differences expressed as percentages across domain-domain/subdomain totals. msFNC connectivity domains include Cerebellar (CB); Visual (VI), comprising Occipitotemporal (OT) and Occipital (OC) subdomains; Paralimbic (PL); Subcortical (SC), comprising Extended Hippocampal (EH), Extended Thalamic (ET), and Basal Ganglia (BG) subdomains; Sensorimotor (SM); Higher Cognition (HC), comprising Insular Temporal (IT), Temporoparietal (TP), and Frontal (FR) subdomains; Triple Network (TN), comprising Central Executive (CE), Default Mode (DM), and Salience (SA) subdomains. Collectively, these results isolate subtype-specific connectivity signatures, emphasizing that each PIN is characterized by unique and reproducible network-level alterations.

fPIN-1 exhibited distinct hyperconnectivity between the cerebellar and sensorimotor domains as well as between subcortical-basal ganglia and triple network-default mode subdomains (Figure 4.A right panel). In contrast, hypoconnectivity emerged between visual-occipitotemporal and sensorimotor domains.

fPIN-2 displayed distinct hyperconnectivity involving dysconnections within subcortical domain, and between subcortical-extended thalamic and visual-occipitotemporal, higher cognition (insular temporal and frontal subdomains) (Figure 4.B right panel). Hypoconnectivity was found within the higher cognition domain and between triple network-central executive, default mode, salience subdomains and subcortical-extended thalamic, basal ganglia subdomains.

mPIN-1 demonstrated pronounced hyperconnectivity between the sensorimotor domain and subcortical subdomains, including extended thalamic and basal ganglia (Figure 4.C right panel). PIN also exhibited hypoconnectivity between the cerebellar and subcortical-basal ganglia, higher cognition-frontal, triple network-salience subdomains (Figure 4.C). Further hypoconnectivity involved within subcortical, as well as within higher cognition domains, between triple network (central executive, default mode subdomains) and sensorimotor, higher cognition-insular temporal subdomain. Figure 4.C right panel highlights these distinct dysconnections reflecting hyperconnective sensorimotor coupling and hypoconnective higher cognition and subcortical connectivity (Figure 4.C right panel).

mPIN-2 showed hyperconnectivity between triple network (central executive and default mode) and higher cognition-temporoparietal with subcortical subdomains (Figure 4.D left and right panels). Hypoconnectivity appeared within subcortical-extended hippocampal and higher cognition-insular temporal subdomains, between higher cognition and sensorimotor domains.

### 2.3. Differential Neurobiology Underlying Phenotypic Measures

Group-level comparisons of the Brief Assessment of Cognition in Schizophrenia composite score (BACS) and symptom measures, such as the Young Mania Rating Scale (YMRS), Montgomery and Asberg Depression Rating Scale (MADRS), and PANSS (includes positive, negative, general, and total scores rated on the Positive and Negative Syndrome Scale^52^) across the four sex-dominant PINs revealed no statistically significant differences (see Supplementary Section.5). We next evaluated whether PIN-specific disrupted subsystems underlie cognitive and symptom outcomes, hypothesizing that PINs exhibit differential levels of disruption within distinct subsystems, which in turn relates to differences in subsystem-associated phenotypic variation across PINs. To test this, we train separate LASSO regression models (a machine learning approach that identifies the most relevant predictors while controlling for overfitting) for each PIN-specific disrupted subsystem (i.e. distinct dysconnectivity features uniquely characterizing that PIN relative to others; Results 2.2.1; Figure 4; See Method 7.5). Each model captures neurocognitive and symptom relationships constrained to a specific disrupted subsystem (see Supplementary Section.6), rather than all msFNC features. These features served as predictors of the BACS composite score, YMRS, MADRS, and PANSS symptom domains. All models were trained on psychosis probands in the discovery set, with L1 regularization and cross-validation used to improve generalizability and reduce overfitting (see Method 7.5), and hyperparameters tuned while adjusting for race, site, sex, and head motion (meanFD). The final models yielded by the discovery set were validated in the hold-out replication set to ensure replicability. All four disrupted subsystems, each characterized by PIN-distinct dysconnectivity profiles, showed significant and replicable associations with BACS composite score (Figure 5.A). Taken together, these findings indicate a possibility that the cognitive impairment noted in PINs may (partially) arise from distinct connectivity subsystem disruptions. Furthermore, the fPIN-2-disrupted subsystem significantly predicted PANSS negative, general, and total symptom scores (see Supplementary Section.6), where no other PIN subsystem showed such associations.

**Figure 5.**
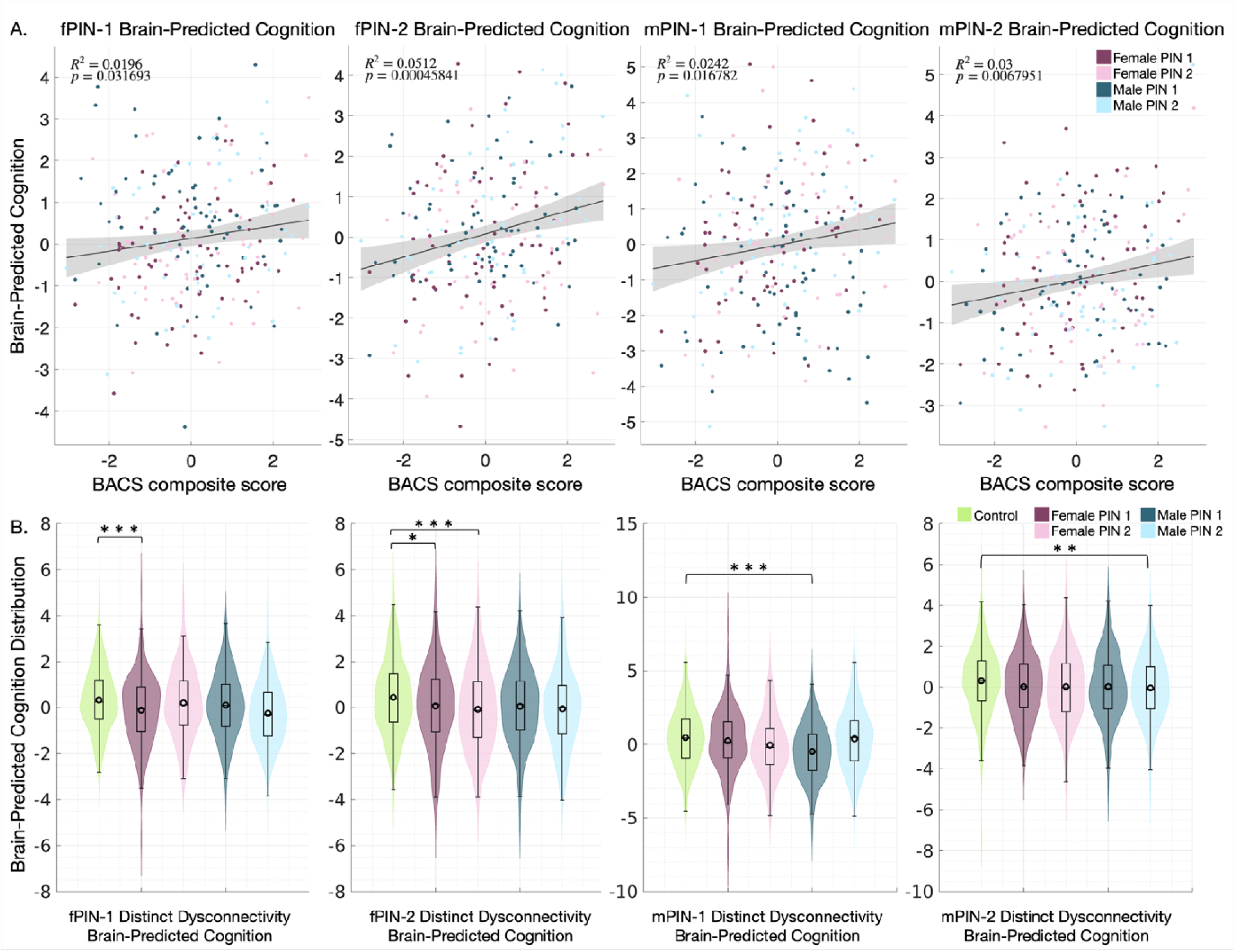
Sex-dominant Psychosis Imaging Neurosubtypes (PINs) Brain-Predicted Cognitive performance (BPC). LASSO regression brain-BACS models from PIN distinct dysconnectivity validating BPC in the replication set. (A) Scatter plots showing the actual Brief Assessment of Cognition in Schizophrenia composite score (BACS) vs. BPC for fPIN-1, fPIN-2, mPIN-1, and mPIN-2, with corresponding R² and p-values. (B) Violin plots show BPC group distributions estimated from different brain-BACS models. Statistical evaluation of BPC between PINs and controls is highlighted (*p < 0.05, **p < 0.01, ** * p < 0.001). These results demonstrate that subtype-specific dysconnectivity patterns carry predictive information about co nitive performance, supporting the functional relevance of sex-dominant PINs.

### 2.4. PIN-Specific Brain-Predicted Cognitive performance

To further verify whether distinct connectivity subsystems underlie cognitive outcomes across PINs, we next examined whether the cognitive performance predicted by the PIN-specific disrupted subsystem, termed as Brain-Predicted Cognitive performance (BPC), also showed PIN-specific deficit compared to controls. BPC is an imaging-derived estimate of cognitive performance derived from the LASSO models (brain-BACS models) developed in the previous section. For each individual, four BPC values were estimated (one from each PIN-specific disrupted subsystem model). We then tested whether predicted cognitive measures would show uniquely significant reductions compared to controls within the corresponding model (i.e., neurosubtype-congruent BPC pattern), reflecting PIN-specific pathways of cognitive deficits, rather than effects from shared neurobiology across subtypes.

For each disrupted subsystem, the corresponding PIN always showed the most control-discriminative power compared to others (Figure 5.B), supporting neurosubtype-congruent relationships and specificity of BPC. Specifically, we observed significant and replicable case–control differences in predicted BACS composite scores for fPIN-1 (t = −3.41, p < 0.001), fPIN-2 (t = −3.49, p < 0.001), mPIN-1 (t = −3.76, p < 0.001), and mPIN-2 (t = −2.68, p < 0.01), when predicted from their respective brain-BACS models. Predicted scores derived from non-matching PIN models did not differ from controls, except when fPIN-1 probands were projected onto fPIN-2’s subsystem, where their predicted scores were modestly lower than controls (t = −2.57, p < 0.05). These findings indicate that although neurocognitive relationships are shared across psychosis probands, the disruption in the subsystem is strongest for the matching PIN, which hence showed the most significant reduction in the predicted BPC compared to controls. Together, these results support the existence of a distinct PIN-specific pathway through which connectivity disruptions impact cognitive performance.

## 3. DISCUSSION

To the best of our knowledge, this study represents the first attempt to identify sex-dominant neurosubtypes and to explicitly account for sex in disentangling the heterogeneity of psychosis. While prior literature has framed sex effects as differences between males and females, our results indicate a more nuanced picture, supporting the notion of heterogeneity both within and across males and females.

In our approach, we addressed sex-related heterogeneity in psychosis using robust multiscale functional connectivity combined with an ensemble bagging framework. Guided by the Neuromark 2.2 template^48^ of 105 multiscale ICNs derived from over 100,000 individuals spanning multiple ICA model orders, we obtained stable, biologically meaningful components that have previously enabled the identification and replication of functionally homogeneous neurosubtypes and cognitive-related biotypes^13,53^. We used the time courses of these ICNs to compute high-dimensional msFNC, from which we extracted low-dimensional embeddings and applied ensemble bootstrapped clustering to derive the sex-dominant PINs.

This approach identified four sex-dominant PINs: two female-dominant (fPIN-1, fPIN-2) and two male-dominant (mPIN-1, mPIN-2). The PINs were not sex-exclusive but demonstrated significant sex overrepresentation and distinct dysconnectivity patterns across major functional domains, supporting sex as a key axis of heterogeneity in psychosis.

These PINs proved robust across two independent checks. First, non-parametric testing confirmed significant sex overrepresentation in three of four PINs in the replication sample (fPIN-1, fPIN-2, mPIN-2); mPIN-1 did not reach significance (p = 0.14), which we attribute to limited replication sample size rather than a lack of validity, given its consistent dysconnectivity patterns and cognitive associations across the discovery and replication sets. Second, the PINs remained consistent following neuroCombat harmonization, supporting robustness to multisite confounding effects rather than an artifact of site variability.

To assess whether this subtype structure was specific to psychosis, we applied the same framework to controls. No control-derived subtype showed significant sex overrepresentation in the replication set (all p > 0.15), indicating that the sex-dominant subtype structure did not replicate among controls; consistent with the PINs reflecting characteristics specific to psychosis heterogeneity rather than a generic outcome of the clustering approach. Given the small replication sample size, and by the same reasoning applied to mPIN-1 above, we interpret this null result cautiously rather than as definitive: larger control cohorts are needed to determine whether comparable sex-related structure might emerge with greater statistical power.

For all four sex-dominant PINs, their disrupted connectivity subsystems, showed replicated neurocognitive associations, indicating that each PIN is characterized by distinct network pathways underlying cognitive impairment. The disrupted system of fPIN-2 extends beyond cognitive performance, showing associations with symptom severity across PANSS negative, general, and total scales. Notably, although there were no statistically significant differences in observed symptom scores and cognitive measures between PINs, BPC revealed PIN-specific patterns, with reduced predicted cognitive performance observed only in the corresponding PIN relative to controls. This suggests clinically similar cognitive impairments may arise from distinct underlying neurobiological disruptions. While prior studies often report better cognitive performance in females^40^, our results did not replicate this pattern, even within individual BACS scores (Supplementary Section 6). Moreover, although sex differences in psychosis are well documented (particularly in SZ), such as earlier onset in males and more affective symptoms in females^54^, such differences have been inconsistent or appear to vary depending on the illness stage. Brand et al. suggest that sex differences between females and males tend to diminish in chronic stages of the disease^55^. Therefore, the inclusion of chronic and clinically stable patients may have further decreased variability, contributing to the absence of group-level clinical differences.

Neuroimaging-derived subtyping is highly sensitive to the imaging modalities and feature representations used, often yielding different or partially overlapping subtype solutions across studies^56^. Given the complexity of psychosis, there may exist different subtyping solutions to capture the heterogeneity of psychosis in distinct dimensions, which are all clinically meaningful^12–24^. In this context, our sex-dominant PINs provide added value: their distinct neurocognitive associations and BPC estimates demonstrate that cognitive impairment is an important axis of sex-related neurobiological heterogeneity in psychosis. Cognitive impairment is one of the core symptomatic dimensions of psychotic disorders, and remains poorly treated despite its strong impact on real-world functioning^57^. Considering that treatment efficacy, dosage, and side-effect profiles vary by sex^55^, our findings suggest that future pharmacological interventions targeting cognitive dysfunction must account for potentially distinct and heterogeneous neurobiological mechanisms both between and within sexes, which we believe has the potential to be clinically meaningful. Males and females should not be treated as two homogeneous biological subgroups, as distinct and heterogeneous neurobiological mechanisms likely underlie cognitive dysfunction within each sex, and one type of treatment is unlikely to fit all.

### 3.1. Sex-Dominant PINs Dysconnectivity

Our study provides new evidence that psychosis is characterized by sex-dominant yet heterogeneous connectivity disruptions. Within each sex, distinct neurosubtypes with more homogeneous functional brain alterations can be identified, suggesting that males and females with psychosis are unlikely to constitute biologically uniform groups. Male neurosubtype mPIN-1 showed extensive dysconnectivity, whereas mPIN-2 showed the least. In contrast, the female neurosubtype fPIN-2 exhibited connectivity disruptions comparable in extent to mPIN-1 but with a distinct pattern.

Previous subtyping studies in psychosis often identify two or three subgroups that differ in symptom severity or the magnitude of biological alterations, leaving open the question of whether such subtypes represent distinct biological entities or simply different severity levels of the same condition^14,17,18,24,53,58^. Our findings suggest biological distinctiveness among the identified sex-dominant PINs. Both female PINs exhibited a comparable number of msFNC alterations (460 vs. 511), but patterns of neurobiological disruptions diverged. Although mPIN-1 had more dysconnectivity than mPIN-2 (748 vs. 272), nearly half of mPIN-2’s alterations were unique (147), reinforcing that it represents a distinct neurobiological subtype rather than a milder form of mPIN-1.

Cerebellar, higher cognition-frontal, and triple network-default mode subdomains msFNC alterations were relatively consistent across two female-dominant PINs (fPIN-1 and fPIN-2), both showing hyperconnectivity. This pattern is consistent with previous reports linking increased activity within these networks in females with schizophrenia to compensatory mechanisms^39,59^ that may mitigate negative symptoms^60^. In contrast, mPIN-1 showed hypoconnectivity, and mPIN-2 more closely resembled female-dominant PINs, indicating that establishing females and males as single biological subgroups may be a misattribution, as a female-specific feature could actually be shared by females and a subgroup of males. Cerebellar-subcortical patterns further reinforce neurosubtype specificity: while past work emphasized hypoconnectivity^61^, we observed hypoconnectivity in fPIN-1 and mPIN-1, hyperconnectivity in fPIN-2, and minimal disruption in mPIN-2. Both female-dominant PINs showed subcortical domain hyperconnectivity (fPIN-2 more than fPIN-1), whereas male-dominant PINs presented hypoconnectivity, patterns also previously linked to negative symptom severity^10,39,62^. Notably, only fPIN-2’s disrupted connectivity subsystem (Supplementary Section.6) is associated with PANSS negative, general, and total scores, driven by cerebellar, subcortical, higher cognition domain, and triple network domains connectivity^25,63–65^. Importantly, relationships previously observed only in males^25^ (subcortical-extended thalamic and triple network-default mode subdomains connectivity association with PANSS) emerge here in fPIN-2, suggesting that biological variability within sexes may have masked such effects in prior sex analyses. Taken together, these results suggest that sex-dominant PINs capture another layer of heterogeneity in psychosis, and a distinct subtype-specific signature suggests multiple neurobiological pathways that may lead to similar clinical syndromes. These sex-dominant PINs provide a framework for sex-stratified biomarker development, with potential to guide personalized treatment and intervention strategies by recognizing distinct neurobiological underpinnings that converge on shared cognitive and clinical outcomes.

### 3.2. Sex-Dominant PINs Differential Cognitive Neurobiology

All four sex-dominant PINs showed that their uniquely disrupted connectivity subsystems were significantly associated with cognitive performance. These predictive features spanned multiple domains (CB, VI, SC, SM, HC, TN), many of which have been consistently identified as major contributors to cognitive performance^10^. Despite these PINs showing comparable group-level cognitive scores, their BPC (derived from PIN-specific distinct dysconnectivity) showed PIN-specific reductions relative to controls, while other PINs showed no significant differences. This suggests that similar cognitive outcomes may arise from differential functional neurobiology across sex-dominant neurosubtypes.

In contrast, previous studies have suggested that neurocognitive relationships are largely shared across sexes in healthy individuals, with substantial overlap in underlying connectivity patterns^66^. Additionally, Damahala et al. reported that psychopathology–brain associations generally do not differ between sexes in children^67^. Our findings were consistent, with all PINs disrupted subsystems being associated with cognitive performance across sex groups. However, PINs have different levels of disruption in these subsystems. Notably, only the fPIN-2 subsystem predicted PANSS general, negative, and total scores, indicating a possibility that the dysconnectivity patterns underlying symptom expression are not all the same across PINs. We posit that protective and risk factors for psychosis, both genetic and environmental, are unequally distributed both between and within males and females^38^. These factors may differentially shape brain development from childhood to adulthood, potentially leading to differential levels of disruptions in dysconnectivity subsystems across PINs, which further link to cognitive impairment and psychopathology. We plan to investigate this further in our future work.

Previous works on the B-SNIP data have also identified cognitive-related subtypes^49^ as well as other characteristic biotypes^15,24^ in psychosis, showing connectivity alterations^53,68^ in the same B-SNIP sample but not accounting for sex differences. Several connectivity features overlap between these biotypes and the current sex-dominant PINs. For instance, Andrés-Camazón et al. reported cerebellar–subcortical hypoconnectivity in cognitive biotype 1, resembling fPIN1 and mPIN1, while cognitive biotype 2 exhibited hyperconnectivity between cerebellar–sensorimotor domains, similar to fPIN1, and hypoconnectivity between sensorimotor-higher cognition domains, akin to mPIN2. Likewise, B-SNIP biotypes^15,68^ showed hyperconnective cerebellar-subcortical, hypoconnective cerebellar and visual-occipital, triple network default-mode subdomain connectivity, patterns expressed by some but not all PINs, since other PINs displayed hyperconnectivity. These findings suggest that connectivity signatures previously identified in psychosis are further refined when sex-related heterogeneity is considered, revealing distinct male and female-dominant neurobiological pathways. The similarity in these brain characteristics underscores the relevance of these networks in neurocognitive relationships and suggests that both subtyping approaches likely capture complementary aspects of this relationship. Some neural features may consistently drive subtyping solutions, whereas others are unique to the adopted approach. Collectively, these observations support the view that the neurobiology of cognitive dysfunction in psychosis is heterogeneous and diverges in sex-related pathways. Importantly, these diverse subtyping solutions may each be clinically meaningful but will require further clinical verification to determine how these differences translate into prognosis, treatment response, or personalized intervention strategies.

## 4. LIMITATIONS

Our study has several limitations: 1) Key confounding factors, such as substance use (e.g., cannabis, nicotine), were not included. 2) B-SNIP probands were clinically stable, which may explain the limited symptom variability and weaker group-level differences. 3) We did not include other neuroimaging modalities, as our focus was specifically on functional network connectivity (FNC). Because neuroimaging-derived subtypes are sensitive across imaging modalities and subtyping features, different imaging choices can yield nonoverlapping subtype solutions, limiting reproducibility and comparisons across studies. 4) FNC is inherently static, relying on full scan temporal correlations among spatially fixed brain networks, which may obscure transient, subject-specific connectivity fluctuations and dynamic spatial reconfigurations known to carry biological relevance^25^ ^70^. 5) The cross-sectional design does not allow inferences about the evolution of sex-specific connectivity patterns.

## 5. FUTURE DIRECTIONS

Key next steps include replication in larger, independent cohorts and incorporation of dynamic connectivity approaches to capture time-varying connectivity patterns^71,72^. Integrating multimodal strategies (structural, functional, and genetic^73^) will be critical for biologically grounded sex-dominant PINs and validating subtype robustness across modalities. Nonlinear representation learning methods, such as deep neural networks that incorporate heterogeneity and covariates, may further refine neurosubtypes^74^. Translationally, longitudinal validation across illness stages and investigation of hormonal and neurotransmitter mechanisms will be essential to determine prognostic value and inform precision-targeted interventions.

## 6. CONCLUSION

We identified four sex-dominant neurosubtypes, two female PINs (fPIN-1, fPIN-2) and two male PINs (mPIN-1 and mPIN-2) using msFNC, uncovering sex-dominant neurobiological heterogeneity. These neurosubtypes were not sex-exclusive but showed male and female overrepresentations, reflecting stratification along the spectrum of male-dominant to female-dominant brain alterations with shared characteristics. Notably, while sex-dominant PINs appear clinically similar, each neurosubtype presented unique brain dysconnectivity further linked to cognitive and symptom outcomes. Particularly, brain Predicted Cognitive performance exhibited PIN-specific reductions relative to controls. This suggests comparable levels of cognitive impairment in neurosubtypes may arise from distinct neurobiological mechanisms, appearing clinically uniform despite heterogeneous neural underpinnings. Notably, the dysconnectivity patterns of female neurosubtype fPIN-2 also showed replicable associations with PANSS negative, general, and total scores. Together, these findings highlight sex as a relevant biological variable in psychosis research and demonstrate that neurosubtypes can be defined by distinct connectivity signatures even when phenotypic measures appear similar. By establishing PIN-specific links between brain networks, cognitive performance, and symptom severity, our study underscores multiscale approaches for developing sex-stratified biomarkers and advancing personalized treatment interventions in psychosis.

## 7. METHODS

### 7.1. Participants

Data used in the study were participants recruited by the B-SNIP consortium 1 and 2^51^. This included controls (*N* = 626) and probands with psychosis (*N* = 1127) meeting the criteria for BP, SZ, and SAD using the DSM-IV Axis I Disorders, Patient Edition (SCID-I/P)^75^, with all participants being unrelated. Inclusion required the availability of rsfMRI, demographic, symptom, and cognitive data. Further details are provided in Supplementary Section 1.

### 7.2. Multiscale Functional Network Connectivity

Imaging data acquisition and preprocessing of rsfMRI are described in Supplementary Section 2. We used the Group ICA of fMRI Toolbox (GIFT) v4.0c package (http://trendscenter.org/software/gift)^71,76^ to perform MOO-ICAR^77^ and generate subject-specific ICNs. The Neuromark 2.2 template (http://trendscenter.org/data) served as the reference in MOO-ICAR, which includes highly replicated 105 ICNs across different spatial scales obtained from over 100k individuals^50,48^. Lastly, we computed subject-level static FNC by calculating pairwise Pearson correlations between the cleaned (Supplementary Section.3) ICN time courses. This process resulted in a 105 × 105 symmetric msFNC matrix for each participant, representing the whole-brain functional connectome (Figure 1, left panel).

### 7.3. Identification of Sex-dominant PINs

#### 7.3.1. PCA on msFNC of Psychosis Probands

The dataset was split into discovery (about 80% of the sample) and replication (about 20% of the sample), with discovery set constructed to preserve a consistent psychosis female-to-male ratio. To ensure that the subtyping was not influenced by age, the discovery msFNC was statistically adjusted for the age covariate using linear regression, and the coefficients applied to the replication set, yielding age-adjusted msFNC for all analyses. PCA was applied to normalized msFNC of probands of the discovery set (i.e., PSY-Discovery), retaining components that accounted for 99% of the variance. These components defined the msFNC subspace of probands. Both PSY-Discovery and PSY-Replication sets msFNC were projected into this subspace (Figure 1, right panel).

#### 7.3.2. Identification of Sex-specific Components and Sex-dominant PINs

Sex differences were assessed in the msFNC subspace using two-sample t-test between female and male proband projections. Components that showed significant sex differences are marked as sex-specific components. K-means clustering separately on these component projections identified two female and male clusters as PINs (Figure 1, right panel). Cluster stability was supported by Calinski-Harabasz index and silhouette based on the sex-specific components to identify subgroups for females and males.

#### 7.3.3. Ensemble Learning to Identify Representative Sex-dominant PINs

To derive robust subtype membership, we implemented an ensemble bagging framework comprising one hundred independent models trained on discovery set with bootstrapping. Each model independently estimated two female and two male clusters. For each model, clustering centroids were identified, and subjects were reassigned to the nearest female/male clusters using a distance-based approach. This reassignment step allowed cross-sex correspondence by matching female and male projections based on closeness to centroids. Cluster labels across models were then aligned using Hungarian algorithm, and the most frequent assignment across models defined each probands representative sex-dominant PIN label. This ensemble approach, aggregating results across models ensured consistent and reproducible neurosubtype identification.

To assess robustness to multisite effects, msFNC was harmonized using neuroCombat. Harmonized data were reassigned to the ensemble models using the same procedure, and the resulting cluster labels were compared with original assignments. Matching between harmonized and original labels was quantified using the Hungarian algorithm and assessed against a null distribution generated from 5000 random permutations of cluster labels.

#### 7.3.4. Sex-dominant PINs Replicability Test

To assess the statistical validity and sex overrepresentation of PINs, we conducted a non-parametric hypothesis test to note their significance in the discovery and replication set (hold-out sample). The null hypothesis assumed that PIN assignment was independent of sex. We generated a null distribution by randomly shuffling sex (female/male) labels 5000 times, while preserving PIN sizes and recalculating sex proportions within each PIN at every iteration. Statistical significance was determined by comparing the observed female-male proportions against the corresponding null distributions, with significance defined using the 95^th^ percentile threshold.

### 7.4. Neurobiological Characterization of Sex-dominant PINs

The neurobiological profile of sex-dominant PINs was characterized by examining msFNC dysconnectivity of each PIN relative to controls, and between PINs. Generalized linear models tested group differences for each msFNC feature, adjusting for race, site, and motion (age was already accounted for). Group analysis results were also then corrected for multiple comparisons using false discovery rate (FDR) correction.

Case-control comparisons above were performed to characterize msFNC dysconnectivity of each sex-dominant PIN. Focusing on the validated msFNC dysconnectivity, i.e., differences observed in the discovery set and further validated in the hold-out replication set, we identified subsystems of unique msFNC dysconnectivity that only showed group differences in individual sex-dominant PINs and defined it as distinct dysconnectivity.

### 7.5. Differential Neurobiology Underlying Phenotypic Measures

To examine whether the PIN-specific disrupted connectivity subsystems contributed to cognitive and symptom variation, we have employed sparse modeling with LASSO regularization. For each sex-dominant PIN, msFNC features from their disrupted subsystem were used as predictors, and separate models were trained across all PSY-Discovery set (Figure 6). Associations with seven phenotypic measures were modelled as outcomes: BACS composite score, MADRS, YMRS, PANSS positive, negative, general, and total (age, race, site, meanFD already accounted for).

**Figure 6.**
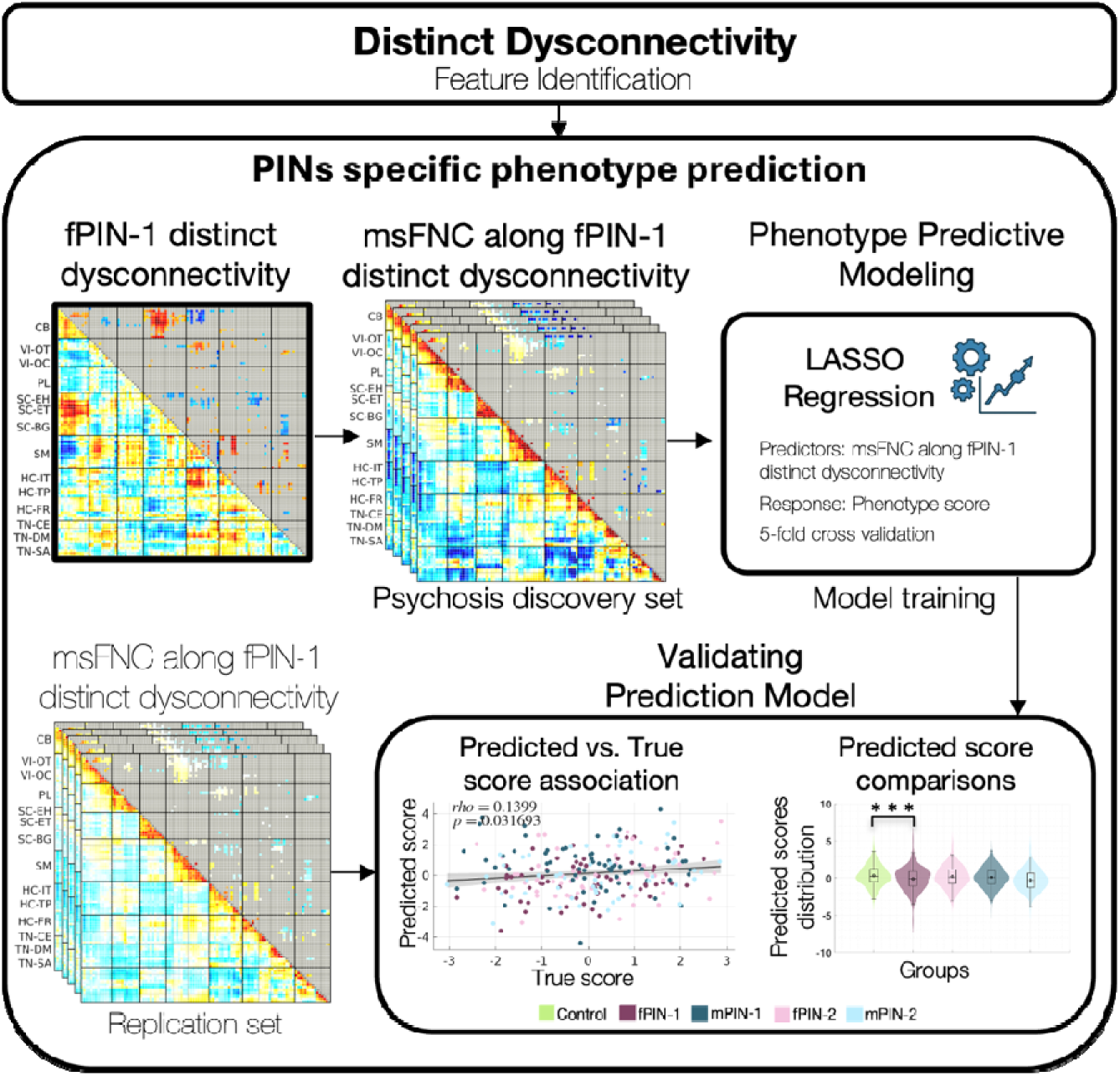
Pipeline for Psychosis Imaging Neurosubtypes (PINs) specific phenotype prediction. Distinct dysconnectivity features defining each PIN-specific disrupted subsystem were first identified and used as predictors in a feature selection step (illustrated here for fPIN-1). Separate LASSO regression models were then trained using these subsystem features from all psychosis probands in the discovery set to predict phenotypic measures (e.g., co nitive performance and symptoms). The trained models were subsequently applied to the independent replication set using the same subsystem features. Model performance was evaluated by assessing the association between predicted and true scores, and their replication consistency. PIN-specificity was further examined by comparing predicted scores across PINs and controls to determine whether each disrupted subsystem captured unique phenotype related variation.

Model training and parameter optimization were conducted exclusively within discovery set PINs, using MATLAB *lassoglm*, with standardized predictors, 5-fold cross-validation, and L1 regularization. To account for stochastic variability in feature selection inherent to LASSO, the entire 5-fold cross-validation was repeated 10 times, yielding different random fold assignments. Model performance was assessed by comparing the statistical relevance of predicted and observed phenotypic measures in the internal validation sets, and the best performing model was selected and applied to the external hold-out replication set to evaluate the consistency of brain-phenotype associations across disrupted subsystems.

PIN-specificity was further assessed by examining whether predicted phenotypic measures differed across PINs within each subsystem specific model. For phenotypic measures available in controls (e.g., BACS), sparse model coefficients were applied to controls’ msFNC data to generate predicted scores, which were then compared between controls and each PIN. For symptom scales relevant to probands (e.g., PANSS, MADRS, and YMRS), PIN-specific differences were evaluated using generalized linear models, with pairwise comparisons between each PIN and all others. PIN specificity was confirmed when a subsystem derived model showed the strongest phenotypic effect for one PIN compared to others.

## 8. DATA AVAILABILITY

B-SNIP 1&2 study data are available through submitting a request with NDA. Any related data that support the findings of this study are available from the corresponding authors upon reasonable request.

## 9. CODE AVAILABILITY

All code used in this study is available from the corresponding authors upon reasonable request.

## 10. ACKNOWLEDGEMENTS

AI has received grant support from the National Institutes of Health (R01MH136665) and Georgia State University’s Research Initiation Grant (RIG) program. VDC has received grant support from the National Institutes of Health (R01MH123610) and the National Science Foundation (2112455). CDC has received funding from Instituto de Salud Carlos III (ISCIII), Spanish Ministry of Science and Innovation (PI23/00625) and the European Commission (grant number 101057182, project Youth-GEMs and grant number 101156514, project YOUTHreach). PAC is supported by the Instituto de Salud Carlos III (ISCIII), Spanish Ministry of Science and Innovation, Río Hortega Program CM24/00125.

## 11. AUTHOR CONTRIBUTION

AI conceived the study and conducted preliminary analyses. RB contributed to study design, performed data preprocessing and completed main analysis, and drafted the manuscript, supplementary materials, and figures, with extensive guidance from JC and AI. PAC contributed to framing the introduction and provided guidance on interpreting results in the discussion. JC and AI provided critical conceptual input and supervision throughout the project, including manuscript revisions. All authors provided feedback on interpretation, contextualization of findings, and manuscript review. The entire manuscript was thoroughly reviewed by all authors, suggesting necessary revisions. All authors have approved the submitted version of the manuscript and agree on the integrity of the work presented.

## 12. COMPETING INTERESTS

We declare that there are no relevant financial and/or non-financial interests in relation to the work submitted.

## Notes

### Competing Interest Statement

The authors have declared no competing interest.

