## Supplementary Materials for "Multiscale Functional Dysconnectivity Reveals Sex-Dominant Neurosubtypes and Predicts Unique Neurocognitive Pathways in Psychosis"

**Supplementary Section.1 Participants**

Bipolar-Schizophrenia Network on Intermediate Phenotypes (B-SNIP) Consortium 1 and 2 (1) recruited participants were used for this study. All participants were fully informed about the study procedure, and written consent was collected. Participants met diagnostic criteria for Bipolar I Disorder with psychotic features (BP), Schizophrenia (SZ), or Schizoaffective Disorder (SAD), using the DSM-IV Axis 1 Disorders, Patient Edition (SCID/P) (3). Data were collected from multiple sites, all following standardized procedures in recruitment, data acquisition, and clinical and cognitive assessments. Broad cognitive evaluation of participants was administered through the Brief Assessment of Cognition in Schizophrenia (BACS) (2). Normative values of these scores stratified across age and sex (2) were used for this study. Clinical interviews rated probands with psychosis on the Young Mania Rating Scale (YMRS). Montgomery-Asberg depression rating scale (MADRS) (4) and Positive and Negative Syndrome Scale (PANSS) (5).

**Supplementary Section.2 Image Acquisition And Preprocessing**

All participants in the study underwent a 5-minute-long resting-state fMRI scan on a 3T MRI scanner. They were instructed to have their eyes open during the scan, focusing on a crosshair displayed on a monitor, while remaining still. Custom build head coil was also employed to minimize head motion during the scan.

Statistical Parametric Mapping toolbox (SPM 12; http://www.fil.ion.ucl.ac.uk/spm/) was used for the scan preprocessing pipeline. The pipeline discarded the initial ten volumes to ensure steady-state magnetization, eliminating unstable signals caused by radio frequency excitations, followed by slice timing correction for temporal differences at each TR using the SPM toolbox. Next steps involved warping scan data to Montreal Neurological Institute (MNI) space using an echo planar imaging template (Calhoun et al., 2017), resampling imaging data to 3 × 3 × 3 mm^3^ voxel space. Spatial smoothing was applied to these images through a Gaussian kernel with a full-width half maximum (FWHM) of 6 mm to enhance the signal-to-noise ratio.

**Supplementary Table 1. Imaging Parameters of B-SNIP1 and B-SNIP2 multisite fMRI data**

| **Site** | **TR (ms)** | **TE (ms)** | **FA (degree)** | **Slice (N)** | **Slice order** | **Acquisition Matrix(mm)** | **Voxel Size (mm)** | **Vendor** |
| --- | --- | --- | --- | --- | --- | --- | --- | --- |
| **B-SNIP1** |  |  |  |  |  |  |  |  |
| Baltimore | 2210 | 30 | 70 | 36 | Interleaved ascending | 64 × 64 | 3.4×3.4×3 | Siemens Triotim |
| Hartford | 1500 | 27 | 70 | 29 | Sequential ascending | 64 × 64 | 3.4×3.4×5 | Siemens Allegra |
| Detroit | 1570 | 22 | 60 | 29 | Sequential ascending | 64 × 64 | 3.4×3.4×4 | Siemens TrioTim |
| Dallas | 1500 | 27 | 60 | 29 | Sequential ascending | 64 × 64 | 3.4×3.4×4 | Philips |
| Chicago | 1775 | 27 | 60 | 29 | Sequential ascending | 64 × 64 | 3.4×3.4×4 | GE Signa HDX |
| **B-SNIP2** |  |  |  |  |  |  |  |  |
| Athens | 2000 | 30 | 60 | 30 | Interleaved ascending | 64×64 | 3.4×3.4×5 | GE HDx |
| Hartford | 2000 | 30 | 60 | 30 | Sequential ascending | 64×64 | 3.4×3.4×5 | Siemens Skyra |
| Boston | 2000 | 30 | 60 | 30 | Sequential ascending | 64×64 | 3.4×3.4×4 | GE HDxt |
| Dallas | 2000 | 30 | 60 | 30 | Sequential ascending | 64×64 | 3.4×3.4×4 | Philips Achieva |
| Chicago | 2000 | 30 | 60 | 30 | Sequential ascending | 64×64 | 3.4×3.4×4 | Philips dStream Achieva |

**Supplementary Section.3 Estimating Subject-Specific Multiscale Functional Network Connectivity**

Subject-specific intrinsic connectivity networks (ICNs) were estimated using Group ICA of fMRI Toolbox (GIFT) v4.0c package with MOO-ICAR, a reference-guided ICA approach that incorporates spatial priors while preserving spatial independence. Spatial priors were obtained from Neuromark 2.2, a standardized multiscale atlas comprising 105 canonical ICNs derived from rsfMRI data of over 100k individuals across multiple studies. Neuromark 2.2 integrates reproducible components identified across ICA model orders (25-200), providing a robust multiscale representation of brain functional organization. Supplementary Table 2 describes Neuromark 2.2 ICN labels, domains, and details and descriptions. Using these spatial priors, subject-specific multiscale ICNs and their corresponding time courses were estimated.

Time courses were subsequently preprocessed by removing linear, quadratic, and cubic trends. Head motion was also accounted for through regression of time courses with motion parameters and their derivatives. Outliers detected based on median absolute deviation and spike threshold (c1) = 2.5 (Jo et al., 2013) were replaced with the best estimate using a third-order spline fit. Finally, preprocessing involved bandpass filtering by applying a fifth-order Butterworth filter with a cutoff frequency of 0.01 Hz – 0.15 Hz. Pearson correlation of these clean, preprocessed 105 time courses results in a 105 × 105 symmetric matrix, estimating the whole-brain multiscale Functional Network Connectivity (msFNC).

**Supplementary Table 2. Neuromark 2.2 reference Domain, Subdomain and Individual ICN Labels**

| **Domain** | **ICN** | **Label** | **Peak Coordinates** | | |
| --- | --- | --- | --- | --- | --- |
|  |  |  | **x** | **y** | **z** |
| **Cerebellar Domain (CB)** | | | | | |
|  | 1 | anterior cerebellum (aCB) | 39 | -40 | -40 |
|  | 2 | posterior cerebellum (pCB) | 30 | -82 | -37 |
|  | 3 | anterior ventromedial cerebellum (avmCB) | 12 | -55 | -52 |
|  | 4 | ventromedial cerebellum (vmCB) | -21 | -55 | -52 |
|  | 5 | right cerebellum (rCB) | 30 | -55 | -43 |
|  | 6 | left cerebellum (lCB) | -27 | -58 | -40 |
|  | 7 | bilateral cerebellum (bCB) | 21 | -67 | -31 |
|  | 8 | medial cerebellum (mCB) | -3 | -61 | -31 |
|  | 9 | anterior dorsomedial cerebellum (admCB) | 0 | -52 | -19 |
|  | 10 | left anterior dorsomedial cerebellum (ladmCB) | -15 | -46 | -25 |
|  | 11 | right anterior dorsomedial cerebellum (radmCB) | 15 | -46 | -22 |
|  | 12 | vermis (Ver) | 0 | -49 | -13 |
|  | 13 | dorsomedial cerebellum (dmCB) | -3 | -64 | -16 |
| **Visual Domain (VI)** | | | | | |
| *Occipitotemporal Subdomain (OT)* | 14 | fusiform gyrus (FG) | 33 | -46 | -16 |
|  | 15 | ventromedial occipitotemporal cortex (vmOTC) | 30 | -49 | -10 |
|  | 16 | left medial occipitotemporal cortex (lmOTC) | -21 | -49 | -7 |
|  | 17 | right medial occipitotemporal cortex (rmOTC) | 21 | -46 | -7 |
|  | 18 | anterior medial occipitotemporal cortex (amOTC) | 15 | -67 | 11 |
|  | 19 | occipitotemporal junction (OTJ) | 45 | -61 | 5 |
| *Occipital Subdomain (OC)* | 20 | extrastriate cortex (ESC) | -42 | -73 | -4 |
|  | 21 | left extrastriate cortex (lESC) | -27 | -76 | -4 |
|  | 22 | right extrastriate cortex (rESC) | 30 | -73 | -1 |
|  | 23 | striate cortex (Str) | -6 | -88 | -4 |
|  | 24 | cuneus (CUN) | -9 | -94 | 26 |
|  | 25 | occipital pole (OP) | 27 | -94 | -4 |
| **Paralimbic Domain (PL)** | | | | | |
|  | 26 | lateral temporal pole (latTP) | 51 | 14 | -25 |
|  | 27 | left temporal pole (lTP) | -30 | 2 | -37 |
|  | 28 | right temporal pole (rTP) | 27 | 5 | -37 |
|  | 29 | left medial temporal cortex (lmTC) | -21 | -7 | -25 |
|  | 30 | left medial temporal pole (lmTP) | -24 | 8 | -25 |
|  | 31 | right medial temporal cortex (rmTC) | 21 | -7 | -25 |
|  | 32 | right medial temporal pole (rmTP) | 27 | 8 | -25 |
|  | 33 | bilateral medial temporal pole (bmTP) | -24 | 5 | -28 |
|  | 34 | bilateral temporal pole (bTP) | 27 | 8 | -34 |
|  | 35 | entorhinal cortex (EC) | 24 | 2 | -34 |
|  | 36 | hippocampal–entorhinal complex (HEC) | 24 | -13 | -19 |
| **Subcortical Domain (SC)** | | | | | |
| *Extended Hippocampal Subdomain (EH)* | 37 | right hippocampus/parahippocampal cortex (rHPC) | 30 | -31 | -4 |
|  | 38 | left hippocampus/parahippocampal cortex (lHPC) | -30 | -34 | -4 |
|  | 39 | thalamus/hippocampus/amygdala (Thal-Amg) | 9 | -31 | 5 |
| *Extended Thalamic Subdomain (ET)* | 40 | thalamus/hippocampus (Thal-Hip) | -3 | -13 | 2 |
|  | 41 | thalamus (Thal) | 9 | -7 | 8 |
|  | 42 | diencephalon/midbrain (DiMid) | 6 | -13 | -4 |
|  | 43 | anterior diencephalon (aDi) | 6 | -4 | -4 |
|  | 44 | right thalamus (rThal) | 21 | -13 | -1 |
|  | 45 | left thalamus (lThal) | -21 | -13 | -1 |
| *Basal Ganglia Subdomain (BG)* | 46 | dorsal striatum/thalamus (DS-Thal) | -21 | 5 | 5 |
|  | 47 | right lentiform nucleus (rLN) | 27 | 5 | -4 |
|  | 48 | left lentiform nucleus (lLN) | -27 | 2 | -4 |
|  | 49 | lentiform nucleus (LN) | 24 | 8 | -1 |
|  | 50 | left caudate (lCaud) | -21 | 17 | 5 |
|  | 51 | right caudate (rCaud) | 21 | 17 | 5 |
|  | 52 | left basal ganglia (lBG) | -18 | 11 | -10 |
|  | 53 | right basal ganglia (rBG) | 18 | 14 | -7 |
|  | 54 | bilateral basal ganglia (bBG) | 9 | 8 | -7 |
| **Sensorimotor Domain (SM)** | | | | | |
|  | 55 | supplementary motor area (SMA) | 0 | -1 | 59 |
|  | 56 | medial supplementary motor area (mSMA) | -9 | 2 | 44 |
|  | 57 | posterior supplementary motor area (pSMA) | -15 | -10 | 62 |
|  | 58 | precentral gyrus (PrCG) | 36 | -10 | 41 |
|  | 59 | superior sensorimotor cortex (sSMC) | 21 | -25 | 59 |
|  | 60 | paracentral lobule (PCL) | 0 | -25 | 65 |
|  | 61 | right superior postcentral gyrus (rsPoCG) | 39 | -22 | 59 |
|  | 62 | left superior postcentral gyrus (lsPoCG) | -39 | -22 | 62 |
|  | 63 | superior parietal lobule (SPL) | 21 | -52 | 71 |
|  | 64 | inferior postcentral gyrus (iPoCG) | -51 | -10 | 32 |
|  | 65 | supramarginal gyrus (SMG) | 54 | -22 | 29 |
|  | 66 | posterior postcentral gyrus (pPoCG) | -54 | -25 | 38 |
|  | 67 | anterior inferior parietal lobe (aIPL) | 60 | -25 | 41 |
|  | 68 | posterior inferior parietal lobe (pIPL) | -60 | -40 | 41 |
| **Higher Cognition Domain (HC)** | | | | | |
| *Insular Temporal Subdomain (IT)* | 69 | left posterior insular cortex (lPIC) | -39 | -7 | 17 |
|  | 70 | right posterior insular cortex (rPIC) | 36 | -13 | 11 |
|  | 71 | dorsal anterior insular cortex (dAIC) | -36 | 5 | 11 |
|  | 72 | ventral anterior insular cortex (vAIC) | -42 | -1 | -10 |
|  | 73 | medial ventral anterior insular cortex (mvAIC) | 42 | 5 | -13 |
|  | 74 | posterior superior temporal gyrus (pSTG) | -39 | -31 | 14 |
|  | 75 | superior temporal gyrus (STG) | 63 | -25 | 5 |
| *Temporoparietal Subdomain (TP)* | 76 | left middle temporal gyrus/temporoparietal junction (lMTG-TPJ) | -48 | -46 | 8 |
|  | 77 | right middle temporal gyrus/temporoparietal junction (rMTG-TPJ) | 48 | -43 | 8 |
|  | 78 | bilateral temporoparietal junction/social mind areas (bTPJ-S) | 57 | -46 | 23 |
|  | 79 | posterior temporal cortex (pTC) | 51 | -46 | 14 |
|  | 80 | inferior posterior temporal cortex (ipTC) | -51 | -46 | -7 |
| *Frontal Subdomain (FR)* | 81 | right inferior frontal gyrus (rIFG) | 51 | 20 | 17 |
|  | 82 | left inferior frontal gyrus (lIFG) | -48 | 26 | 5 |
|  | 83 | ventrolateral prefrontal cortex (VLPFC) | 51 | 23 | 17 |
|  | 84 | inferior left ventrolateral prefrontal cortex (ilVLPFC) | -48 | 20 | 23 |
|  | 85 | left ventrolateral prefrontal cortex (lVLPFC) | -45 | 20 | 23 |
|  | 86 | posterior inferior frontal gyrus (pIFG) | -36 | 14 | 29 |
|  | 87 | orbitofrontal cortex (OFC) | -24 | 50 | -7 |
|  | 88 | frontal pole (FP) | -30 | 62 | 8 |
|  | 89 | Frontal eye field area (FEF) | -15 | 11 | 53 |
|  | 90 | anterior supplemental motor area (aSMA) | -9 | 8 | 62 |
| **Triple Network Domain (TN)** | | | | | |
| *Central Executive Network Subdomain (CE)* | 91 | right inferior parietal lobe/dorsolateral prefrontal cortex (rIPL-rDLPFC) | 48 | -58 | 47 |
|  | 92 | left inferior parietal lobe/dorsolateral prefrontal cortex (lIPL-lDLPFC) | -45 | -61 | 50 |
|  | 93 | bilateral inferior parietal lobe/dorsolateral prefrontal cortex (bIPL-lDLPFC) | -51 | -58 | 41 |
| *Default Network Subdomain (DN)* | 94 | right inferior parietal lobe/posterior cingulate cortex (rIPL-rPCC) | 48 | -64 | 38 |
|  | 95 | left inferior parietal lobe/posterior cingulate cortex (lIPL-lPCC) | -48 | -64 | 35 |
|  | 96 | posterior cingulate cortex (PCC) | -12 | -58 | 20 |
|  | 97 | ventral posterior cingulate cortex (vPCC) | -12 | -52 | 11 |
|  | 98 | ventral precuneus (vPRCU) | -9 | -67 | 35 |
|  | 99 | dorsal precuneus (dPRCU) | -6 | -55 | 50 |
|  | 100 | dorsomedial prefrontal cortex (dmPFC) | 0 | 47 | 35 |
|  | 101 | medial prefrontal cortex (mPFC) | 0 | 50 | 20 |
| *Salience Network Subdomain (SN)* | 102 | anterior cingulate cortex (ACC) | 0 | 41 | 2 |
|  | 103 | dorsal anterior cingulate cortex (dACC) | 0 | 35 | 11 |
|  | 104 | anterior cingulate cortex/anterior insula (ACCAI) | 0 | 32 | 20 |
|  | 105 | anterior insula/anterior cingulate cortex (AIACC) | 33 | 23 | -4 |

**Supplementary Section.4 Sex-Dominant PINs msFNC Characterization**

Pairwise msFNC comparisons between controls and each PIN, as well as between PINs, are represented in Supplementary Figure 1. Figures.1A-1J show statistically significant connectivity differences replicated across discovery and replication sets, while Figures.1K-1T. summarize replication set results. The following sections describe msFNC characteristics of each PIN, highlighting domain and subdomain level hyperconnectivity, hypoconnectivity of PINs relative to controls, and connectivity differences between PINs.

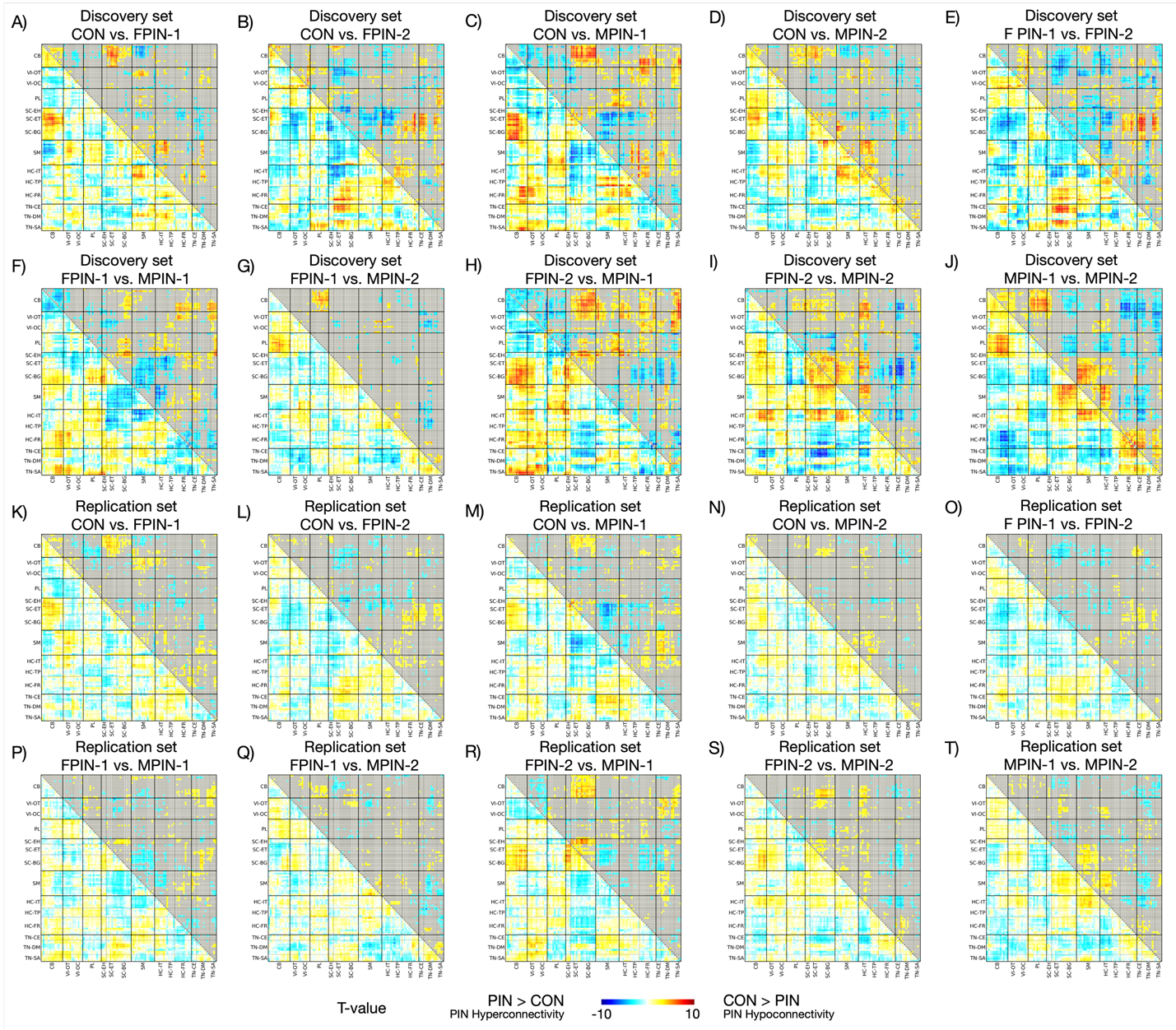

**Supplementary Figure.1: Group Differences: Controls (CON) vs. Sex-Dominant Psychosis Imaging Neurosubtypes (PINs), and between PINs along Multiscale Functional Network Connectivity (msFNC) in Discovery and Replication sets.** msFNC represents connectivity between 105 Multiscale Intrinsic Connectivity Networks (msICNs). Refer to Main Figure.2 and Supplementary Table 1 for additional information on the 105 multiscale intrinsic connectivity networks. 105 × 105 matrices display the t-value in the lower triangle, where each cell reflects a group difference in connectivity between two msICNs, estimated using generalized linear models. The upper triangle shows the statistical significance using -log(q-value).*sign(t-value) for the discovery set and -log(p-value).*sign(t-value) for the replication set, with non-significant msFNC pairs grayed out. The t-value ranges from -10 to 10. A-D) msFNC group difference matrix between CON vs. PINs in discovery and replication sets and common group differences between them. A) msFNC group difference matrix between CON vs. fPIN-1. B) msFNC group difference matrix between CON vs. fPIN-2. C) msFNC group difference matrix between CON vs. mPIN-1. D) msFNC group difference matrix between CON vs. mPIN-2. E-J) msFNC group difference matrix between PINs in the discovery and replication sets and common differences between them. E) msFNC group difference matrix between fPIN-1 vs. fPIN-2. F) msFNC group difference matrix between fPIN-1 vs. mPIN-1. G) msFNC group difference matrix between fPIN-1 vs. mPIN-2. H) msFNC group difference matrix between fPIN-2 vs. mPIN-1. I) msFNC group difference matrix between fPIN-2 vs. mPIN-2. J) msFNC group difference matrix between mPIN-1 vs. mPIN-2. K-N) msFNC group difference matrix between CON vs. PINs in replication set K) msFNC group difference matrix between CON vs. fPIN-1. L) msFNC group difference matrix between CON vs. fPIN-2. M) msFNC group difference matrix between CON vs. mPIN-1. N) msFNC group difference matrix between CON vs. mPIN-2. O-T) msFNC group difference matrix between PINs in replication set. O) msFNC group difference matrix between fPIN-1 vs. fPIN-2. P) msFNC group difference matrix between fPIN-1 vs. mPIN-1. Q) msFNC group difference matrix between fPIN-1 vs. mPIN-2. R) msFNC group difference matrix between fPIN-2 vs. mPIN-1. S) msFNC group difference matrix between fPIN-2 vs. mPIN-2. T) msFNC group difference matrix between mPIN-1 vs. mPIN-2.

***4.1. Differences within functional domains.***

In the cerebellar domain, fPIN-1, fPIN-2 and mPIN-2 exhibited hypoconnectivity (Supplementary Figure.1A, 1B and 1D), while mPIN-1 showed no connectivity difference when compared with controls (Supplementary Figure.1C). Comparing the PINs among themselves, mPIN-1 and fPIN-2 had increased cerebellar domain connectivity than fPIN-1 (Supplementary Figure.1F and 1E), while all other neurosubtypes comparison with each other show no connectivity difference (Supplementary Figure.1E, 1H-J). Within the subcortical domain, fPIN-2 showed hyperconnectivity compared to controls along all subcortical subdomains (extended hippocampal, extended thalamic, basal ganglia; Supplementary Figure 1B); however, fPIN-1 presented hyperconnectivity between subcortical-extended hippocampal and subcortical-extended thalamic and basal ganglia subdomains (Supplementary Figure 1A). On the contrary, male PINs presented hypoconnectivity relative to controls (Supplementary Figure 1C and 1D). Examining the PINs among themselves, fPIN-2 had increased connectivity within the subcortical domain relative to all other neurosubtypes (Supplementary Figure 1E, 1H and 1I), while mPIN-1 had higher subcortical-extended hippocampal and subcortical-extended thalamic and basal ganglia subdomains connectivity (Supplementary Figure 1F and 1H). In the higher cognition domain, all PINs showed hypoconnectivity relative to controls (Supplementary Figure 1A-D), however, mPIN-1 also exhibited hyperconnectivity between higher cognition-frontal and temporoparietal subdomain connectivity (Supplementary Figure 1C), and mPIN-2 showed hyperconnectivity between higher cognition-frontal and insular temporal subdomain (Supplementary Figure 1D). mPIN-1 had increased connectivity compared to mPIN-2 within higher cognition-frontal and insular temporal subdomain connectivity, while mPIN-2 had increased connectivity compared to mPIN-1 between higher cognition-frontal and insular temporal subdomain connectivity (Supplementary Figure 1J). fPIN-1 showed decreased connectivity compared to mPIN-1 along higher cognition-frontal subdomain connectivity (Supplementary Figure.1F). fPIN-2 had increased connectivity relative to mPIN-2 between the higher cognition-insular temporal subdomains (Supplementary Figure.1I).

***4.2. Differences between functional domains***

**Cerebellar – Subcortical Domains Connectivity:** PINs except fPIN-2 were overall hypoconnective compared to controls (Supplementary Figure 1A-D). fPIN-1 showed hypoconnectivity, while fPIN-2 showed hyperconnectivity relative to controls within the cerebellar and subcortical domain connectivity (Supplementary Figure 1A and 1B). Comparing male neurosubtypes with controls, both mPIN-1 and mPIN-2 (with less effect) showed hypoconnectivity relative to controls across cerebellar and all subdomains of subcortical connectivity (Supplementary Figure 1C and 1D). PINs along cerebellar and subcortical domains: fPIN-2 had increased connectivity compared to fPIN-1 (Supplementary Figure 1E). mPIN-1 also had increased connectivity relative to mPIN-2 (Supplementary Figure.1J). fPIN-2 comparison to male neurosubtypes revealed that the female neurosubtype had increased connectivity compared to either of them within cerebellar-subcortical connectivity (Supplementary Figure 1F-I).

**Cerebellar – Sensorimotor Domains Connectivity:** Both female neurosubtypes, fPIN-1 and fPIN-2, had hyperconnectivity compared to controls in cerebellar and sensorimotor domains connectivity (Supplementary Figure 1A and 1B). Female neurosubtype comparison showed that fPIN-1 had increased connectivity compared to fPIN-2 (Supplementary Figure 1E). Similarly, mPIN-2 had increased connectivity compared to mPIN-1 (Supplementary Figure 1J). Comparing female and male neurosubtypes, fPIN-1 had increased connectivity compared to mPIN-1 (Supplementary Figure 1F) but relative to mPIN-2 did not show any significant difference in connectivity along the cerebellar-sensorimotor domains (Supplementary Figure 1G).

**Visual – Subcortical Domains Connectivity:** fPIN-2 and mPIN-1 showed hyperconnectivity relative to controls along the visual-subcortical domains connectivity (Supplementary Figure.1B and 1C), while mPIN-2 and controls did not show any significant connectivity difference (Supplementary Figure.1D). PINs comparisons between each other showed fPIN-2 had higher connectivity than fPIN-1, mPIN-1 and mPIN-2 (Supplementary Figure.1E, 1H-I) between visual-subcortical domains, while male neurosubtypes had no connectivity difference (Supplementary Figure.1J).

**Subcortical – Sensorimotor Domains Connectivity:** Within the differences of subcortical-sensorimotor domain connectivity between PINs and controls, PINs overall showed hyperconnectivity relative to controls (Supplementary Figure 1A-D). fPIN-1, fPIN-2 and mPIN-2 were hyperconnected relative to controls along the subcortical-extended hippocampal, extended thalamic subdomain connectivity with the sensorimotor domain (Supplementary Figure 1A, 1B and 1D). While mPIN-1 showed hyperconnectivity relative to controls, the difference was across complete subcortical (extended thalamic, extended thalamic and basal ganglia subdomains) connectivity with the sensorimotor domain (Supplementary Figure 1C). Female neurosubtypes had little difference, with fPIN-2 presenting slightly higher connectivity than fPIN-1 (Supplementary Figure 1E) among fewer ICNs of subcortical-sensorimotor domains connectivity. However, mPIN-1 had increased connectivity compared to fPIN-1 and mPIN-2 (Supplementary Figure 1F and 4J).

**Higher Cognition – Cerebellar Domains Connectivity:** fPIN-1 and fPIN-2 were hyperconnected relative to controls along higher cognition-insular temporal subdomain connectivity with cerebellar (Supplementary Figure 1A-B). But comparison of male neurosubtype with controls, mPIN-1 showed hypoconnectivity relative to controls along the complete higher cognition domain (insular temporal, temporoparietal and frontal subdomains) connectivity with cerebellar (Supplementary Figure 1C). Between female neurosubtypes, fPIN-2 had higher cerebellar and higher cognition-insular temporal subdomain connectivity than fPIN-1 (Supplementary Figure 1E), while mPIN-2 had higher cerebellar and higher cognition-frontal subdomain connectivity relative to mPIN-1 (Supplementary Figure 1J). Between sexes, fPIN-2 had higher connectivity than mPIN-2 in the cerebellum, and higher cognition-insular temporal subdomain connectivity (Supplementary Figure 1I), but fPIN-1 had higher cerebellar and higher cognition-frontal subdomain connectivity than mPIN-1 (Supplementary Figure 1F).

**Higher Cognition – Subcortical Domains Connectivity:** PINs were hyperconnective relative to controls along the higher cognition-subcortical domain connectivity (Supplementary Figure 1A-D), however, the difference was noted between subdomains. fPIN-1 and mPIN-2 showed this significant difference along the complete higher cognition with subcortical-extended hippocampal, extended thalamic subdomains (Supplementary Figure 1A and 1D), while fPIN-2 showed hypoconnectivity compared to controls along subcortical and higher cognition-frontal subdomains (Supplementary Figure 1B). fPIN-1 showed increased connectivity in subcortical and higher cognition-insular temporal subdomain connectivity compared to fPIN-2, while showing reduced connectivity between subcortical and higher cognition-frontal subdomain connectivity (Supplementary Figure 1E). mPIN-1 relative to mPIN-2 had increased connectivity between higher cognition-frontal and subcortical but reduced connectivity between higher cognition-insular temporal and subcortical domains (Supplementary Figure 1J). Between sexes, mPIN-1 had higher connectivity than fPIN-2 along subcortical and higher cognition-frontal subdomain connectivity (Supplementary Figure 1H)

**Higher Cognition – Sensorimotor Domains Connectivity:** fPIN-1 and mPIN-2 showed hypoconnectivity relative to controls between sensorimotor and higher cognition-temporoparietal, FN subdomain connectivity (Supplementary Figure 1A and 1D). However, mPIN-1 showed hypoconnectivity relative to controls between sensorimotor and higher cognition-insular temporal subdomain connectivity (Supplementary Figure 1C). Notably, fPIN-1 and mPIN-2 had no connectivity difference between each other (Supplementary Figure 1G), and similarly, fPIN-2 and mPIN-1 also did not present any connectivity difference (Supplementary Figure 1H). However, fPIN-1 had reduced connectivity compared to fPIN-2, and mPIN-2 also had reduced connectivity compared to mPIN-1 (Supplementary Figure 1J) between higher cognition-sensorimotor domain connectivity.

**Triple Networks – Cerebellar Domains Connectivity:** PINs did not reveal connectivity differences relative to controls, except where mPIN-1 showed hypoconnectivity (Supplementary Figure 1C), and mPIN-2 showed hyperconnectivity (Supplementary Figure 1D) relative to controls in cerebellar and triple network domain connectivity. mPIN-1 had higher connectivity relative to fPIN-1, fPIN-2 and mPIN-2 along the cerebellar-triple network domains connectivity (Supplementary Figure.1F, 1H-I). However, fPIN-1 had higher connectivity than fPIN-2 in cerebellar-triple network domains connectivity (Supplementary Figure 1E).

**Triple Networks – Subcortical Domains Connectivity: The** fPIN-1 group showed hypoconnectivity, and fPIN-1 showed hyperconnectivity, relative to controls along the subcortical-extended thalamic and triple network-salience subdomains connectivity (Supplementary Figure 1A-B). Comparing the PINs among themselves, a notable difference was observed between fPIN-1 and fPIN-2, where fPIN-1 showed higher connectivity relative to fPIN-2 (Supplementary Figure 1E) along the subcortical-triple network domain connectivity. Male neurosubtypes mPIN-1 and mPIN-2 had very few differences between each other, where mPIN-2 had higher connectivity (Supplementary Figure 1J). However, fPIN-2 had reduced connectivity compared to mPIN-1 and mPIN-2 along subcortical and triple network domain connectivity (Supplementary Figure 1H-I).

**Triple Networks – Sensorimotor Domains Connectivity:** mPIN-1 and fPIN-1 showed hypoconnectivity among sensorimotor and triple network-central executive network, default network subdomains connectivity (Supplementary Figure 1C and 4A), while mPIN-2 and fPIN-2 had no differences. Within the PINs comparisons, fPIN-1 had slightly increased connectivity compared to mPIN-1 among ICNs of sensorimotor-triple network domains connectivity (Supplementary Figure.1F); however, only male neurosubtype differences were notable, where mPIN-1 showed reduced connectivity relative to mPIN-2 along sensorimotor and triple network-central executive network, salience subdomains connectivity (Supplementary Figure 1J).

**Triple Networks – Higher Cognition Domains Connectivity:** mPIN-1 and fPIN-2 revealed hypoconnectivity relative to controls, along the higher cognition-insular temporal subdomain connectivity with triple network domains (Supplementary Figure 1C), while mPIN-2 revealed hyperconnectivity (Supplementary Figure 1D). fPIN-1 also showed hypoconnectivity relative to controls but along the higher cognition-temporoparietal, frontal subdomain connectivity with triple network-central executive network, salience subdomains (Supplementary Figure 1B). Female neurosubtypes revealed few dysconnectivity among each other along the higher cognition-triple network domains connectivity (Supplementary Figure 1E). fPIN-2 and mPIN-1 both had reduced connectivity along higher cognition-insular temporal triple network domains connectivity, but had higher connectivity in higher cognition-frontal and triple network domain connectivity, relative to mPIN-2 (Supplementary Figure 1I and 1J).

**Supplementary Section 5. Sex-Dominant PINs Clinical Characterization**

**Supplementary Table 3. Relationship between Sex-Dominant PINs and DSM Diagnoses**

| **Sex-dominant PIN vs. DSM Diagnoses** | | **Sex-dominant PIN** | | | |
| --- | --- | --- | --- | --- | --- |
|  |  | fPIN-1 | fPIN-2 | mPIN-1 | mPIN-2 |
| **DSM Diagnoses Discovery Set** | SZ | 83 | 77 | 114 | 82 |
|  | BP | 61 | 63 | 63 | 60 |
|  | SAD | 66 | 81 | 69 | 72 |
|  | $\chi^{2}$ | 7.54 | | | |
|  | P-value | 0.27 | | | |
| **DSM Diagnoses Replication Set** | SZ | 24 | 18 | 33 | 15 |
|  | BP | 14 | 15 | 14 | 16 |
|  | SAD | 27 | 23 | 18 | 19 |
|  | $\chi^{2}$ | 7.97 | | | |
|  | P-value | 0.24 | | | |

In both discovery and replication sets, sex-dominant PIN membership was not associated with DSM diagnoses (Supplementary Table 3). Although diagnoses were unevenly distributed across PINs, all diagnostic categories were represented in each neurosubtype. Notably, mPIN-1 showed a higher proportion of schizophrenia in both datasets. Age of onset distributions were also examined across sex-dominant PINs, as well as separately for female and male probands. Distribution patterns were modeled using Gaussian mixture models with one and two components (best reported in Supplementary Figure 2) to assess distributional patterns.

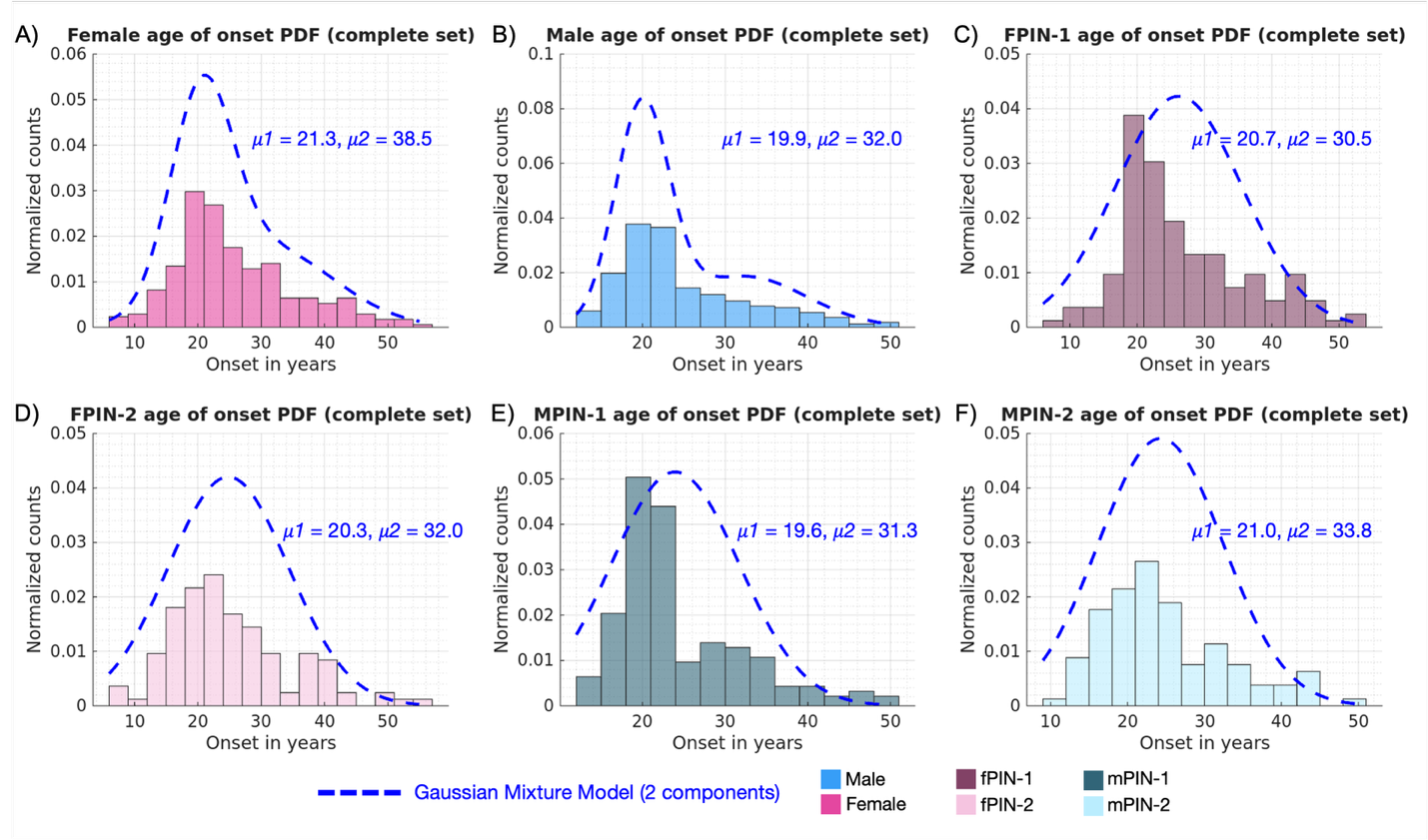

**Supplementary Figure.2: Sex-dominant PINs Age of Onset Distributions.** A-B)Age of onset of the complete set; both Discovery and Replication sets; both female and male proband onset distributions are in part A and B. A) Female proband age of onset distribution. B) Male proband age of onset distribution. C-F) Age of onset distributions of sex-dominant PINs. C) fPIN-1 age of onset distribution. D) fPIN-2 age of onset distribution. E) mPIN-1 age of onset distribution. F) mPIN-2 age of onset distribution.

Additional clinical characterization examined group differences in YMRS, MADRS, PANSS (positive, negative, general, and total), BACS composite score, and age of onset across discovery, replication, and combined datasets. Generalized linear models were used to compare females vs. males, and between PINs while statistically adjusting for age, race, and site. P-values were corrected for multiple comparisons using the False Discovery Rate method. Results are summarized in Supplementary Table 4. No statistically significant differences were observed between PINs for any clinical measure in either the discovery or replication set, or in the combined dataset.

**Supplementary Table 4. Group Differences between Sex-Dominant PINs along Clinical Ratings**

| **Clinical Ratings** | **Discovery set** | | **Replication set** | | **Complete set** | |
| --- | --- | --- | --- | --- | --- | --- |
|  | **T-value** | **P-value (FDR)** | **T-value** | **P-value** | **T-value** | **P-value (FDR)** |
| *Young Mania Rating Scale (YMRS)* |  |  |  |  |  |  |
| female vs. male | 1.49 | 0.77 | -1.50 | 0.13 | 0.69 | 0.66 |
| fPIN-1 vs. fPIN-2 | 0.15 | 0.93 | 1.15 | 0.25 | 0.62 | 0.68 |
| fPIN-1 vs. mPIN-1 | 1.12 | 0.77 | -0.02 | 0.98 | 1.38 | 0.44 |
| fPIN-1 vs.mPIN-2 | 1.55 | 0.77 | 1.96 | 0.05 | 2.32 | 0.24 |
| fPIN-2 vs. mPIN-1 | 1.04 | 0.81 | 0.28 | 0.78 | 1.03 | 0.52 |
| fPIN-2 vs.mPIN-2 | 1.13 | 0.77 | 1.87 | 0.06 | 1.68 | 0.33 |
| mPIN-1 vs.mPIN-2 | 0.59 | 0.86 | 1.72 | 0.09 | 1.17 | 0.49 |
| *Montgomery and Asberg Depression Rating Scale (MADRS)* |  |  |  |  |  |  |
| female vs. male | 4.34 | < 0.001 | 2.21 | 0.03 | 4.82 | < 0.001 |
| fPIN-1 vs. fPIN-2 | 0.23 | 0.93 | -0.70 | 0.48 | -0.24 | 0.85 |
| fPIN-1 vs. mPIN-1 | 2.00 | 0.47 | 0.15 | 0.88 | 1.94 | 0.29 |
| fPIN-1 vs.mPIN-2 | 0.62 | 0.86 | 1.15 | 0.25 | 1.01 | 0.52 |
| fPIN-2 vs. mPIN-1 | 1.24 | 0.77 | 2.00 | 0.05 | 1.95 | 0.29 |
| fPIN-2 vs.mPIN-2 | 0.58 | 0.86 | 1.67 | 0.10 | 1.17 | 0.49 |
| mPIN-1 vs.mPIN-2 | -1.36 | 0.77 | 1.27 | 0.21 | -0.84 | 0.60 |
| *Positive and Negative Syndrome Scale (PANSS) positive* |  |  |  |  |  |  |
| female vs. male | 0.24 | 0.93 | -2.51 | 0.01 | -0.88 | 0.59 |
| fPIN-1 vs. fPIN-2 | 0.05 | 0.98 | 0.77 | 0.44 | 0.53 | 0.72 |
| fPIN-1 vs. mPIN-1 | 1.42 | 0.77 | 1.24 | 0.22 | 2.10 | 0.29 |
| fPIN-1 vs.mPIN-2 | 1.20 | 0.77 | 1.26 | 0.21 | 1.58 | 0.36 |
| fPIN-2 vs. mPIN-1 | 1.37 | 0.77 | 1.24 | 0.22 | 1.75 | 0.32 |
| fPIN-2 vs.mPIN-2 | 0.49 | 0.92 | 0.80 | 0.43 | 0.84 | 0.60 |
| mPIN-1 vs.mPIN-2 | -0.21 | 0.93 | -0.12 | 0.91 | -0.24 | 0.85 |
| *PANSS negative* |  |  |  |  |  |  |
| female vs. male | -3.13 | 0.03 | -2.20 | 0.03 | -3.92 | < 0.01 |
| fPIN-1 vs. fPIN-2 | 0.22 | 0.93 | -1.66 | 0.10 | -0.18 | 0.88 |
| fPIN-1 vs. mPIN-1 | 0.25 | 0.93 | 0.24 | 0.81 | 0.49 | 0.72 |
| fPIN-1 vs.mPIN-2 | 1.51 | 0.77 | 0.97 | 0.33 | 1.81 | 0.31 |
| fPIN-2 vs. mPIN-1 | -0.28 | 0.93 | 1.45 | 0.15 | 0.63 | 0.68 |
| fPIN-2 vs.mPIN-2 | 0.48 | 0.92 | 2.25 | 0.03 | 1.54 | 0.36 |
| mPIN-1 vs.mPIN-2 | 1.01 | 0.81 | 0.56 | 0.58 | 1.05 | 0.51 |
| *PANSS general* |  |  |  |  |  |  |
| female vs. male | 1.35 | 0.77 | -0.43 | 0.67 | 0.90 | 0.59 |
| fPIN-1 vs. fPIN-2 | -0.63 | 0.86 | -1.34 | 0.18 | -1.07 | 0.51 |
| fPIN-1 vs. mPIN-1 | 0.82 | 0.86 | 0.46 | 0.64 | 1.17 | 0.49 |
| fPIN-1 vs.mPIN-2 | 0.73 | 0.86 | 1.69 | 0.09 | 1.40 | 0.44 |
| fPIN-2 vs. mPIN-1 | 0.58 | 0.86 | 2.42 | 0.02 | 1.79 | 0.31 |
| fPIN-2 vs.mPIN-2 | 0.78 | 0.86 | 3.25 | 0.00 | 2.03 | 0.29 |
| mPIN-1 vs.mPIN-2 | 0.14 | 0.93 | 1.22 | 0.22 | 0.47 | 0.72 |
| *PANSS total* |  |  |  |  |  |  |
| female vs. male | -0.26 | 0.93 | -1.74 | 0.08 | -1.12 | 0.49 |
| fPIN-1 vs. fPIN-2 | -0.24 | 0.93 | -0.96 | 0.34 | -0.43 | 0.74 |
| fPIN-1 vs. mPIN-1 | 0.95 | 0.84 | 0.71 | 0.48 | 1.43 | 0.43 |
| fPIN-1 vs.mPIN-2 | 1.23 | 0.77 | 1.55 | 0.12 | 1.78 | 0.31 |
| fPIN-2 vs. mPIN-1 | 0.64 | 0.86 | 2.11 | 0.04 | 1.67 | 0.33 |
| fPIN-2 vs.mPIN-2 | 0.72 | 0.86 | 2.63 | 0.01 | 1.80 | 0.31 |
| mPIN-1 vs.mPIN-2 | 0.34 | 0.93 | 0.76 | 0.45 | 0.51 | 0.72 |
| *Brief Assessment of Cognition in Schizophrenia - composite score (BACS-COMP)* |  |  |  |  |  |  |
| female vs. male | -1.57 | 0.31 | 0.41 | 0.68 | -1.19 | 0.49 |
| fPIN-1 vs. fPIN-2 | 0.11 | 0.93 | -0.57 | 0.57 | -0.30 | 0.88 |
| fPIN-1 vs. mPIN-1 | -1.78 | 0.22 | -0.18 | 0.85 | -1.87 | 0.20 |
| fPIN-1 vs.mPIN-2 | -2.64 | 0.07 | -0.28 | 0.78 | -2.73 | < 0.05 |
| fPIN-2 vs. mPIN-1 | -1.35 | 0.38 | 0.63 | 0.53 | -0.89 | 0.69 |
| fPIN-2 vs.mPIN-2 | -1.92 | 0.18 | 0.38 | 0.71 | -1.75 | 0.24 |
| mPIN-1 vs.mPIN-2 | -0.40 | 0.85 | -0.01 | 0.99 | -0.43 | 0.81 |
| *Age of Onset* |  |  |  |  |  |  |
| female vs. male | 1.99 | 0.25 | -0.11 | 0.91 | 1.51 | 0.32 |
| fPIN-1 vs. fPIN-2 | 1.35 | 0.50 | -0.50 | 0.62 | 0.99 | 0.62 |
| fPIN-1 vs. mPIN-1 | 1.68 | 0.36 | 1.71 | 0.09 | 1.90 | 0.30 |
| fPIN-1 vs.mPIN-2 | 1.89 | 0.26 | 1.31 | 0.20 | 1.75 | 0.30 |
| fPIN-2 vs. mPIN-1 | 0.33 | 0.87 | 0.36 | 0.72 | 0.63 | 0.77 |
| fPIN-2 vs.mPIN-2 | -0.15 | 0.93 | 1.53 | 0.13 | 0.24 | 0.85 |
| mPIN-1 vs.mPIN-2 | -0.18 | 0.93 | 0.75 | 0.46 | -0.29 | 0.85 |

We further evaluated whether medication usage influenced PINs identification by testing whether specific neurosubtypes were associated with medication classes. Medication data from B-SNIP-1 (N=471 probands with psychosis) were analyzed, and medications included psychotropics, antipsychotics, antidepressants, mood stabilizers, anxiolytics, anticonvulsants, antihistamines, and stimulants. Chi-squared tests of independence were performed for each medication class, with p-values corrected for multiple comparisons. No significant associations were observed between PIN membership and medication use, indicating that PIN identification was not driven by medication effects (Supplementary Table 5).

**Supplementary Table 5. Chi-square tests of independence between sex-dominant PIN membership and medication use**

| Medications | $\chi^{2}$ | P-value (FDR) |
| --- | --- | --- |
| N = 471 |  |  |
| Psychotropic | 6.33 | 0.15 |
| Antipsychotic primary | 7.45 | 0.12 |
| Antidepressant primary | 0.73 | 0.87 |
| Moodstabilizer primary | 2.50 | 0.54 |
| Anxiolytic primary | 0.06 | 0.12 |
| Anticonvulsant primary | 0.04 | 0.12 |
| Antihistamine primary | 0.05 | 0.12 |
| Stimulant primary | 0.28 | 0.38 |

**Supplementary Section.6 Differential Neurobiology Underlying Phenotypic Measures**

All four PIN-specific disrupted subsystems showed significant and replicable associations with BACS composite score (Main Results Section 2.3). Additionally, fPIN-2’s disrupted subsystem significantly predicted PANSS negative, general, and total scores (Supplementary Figure 3). Predicted PANSS negative scores were higher for fPIN-2 than fPIN-1 (t = 2.72, p < 0.01), mPIN-1 (t = 2.60, p < 0.05), and mPIN-2 (t = 2.19, p < 0.05). Predicted PANSS general scores were also higher for fPIN-2 than fPIN-1 (t = 2.27, p < 0.05) and mPIN-1 (t = 3.55, p < 0.001). Similarly, predicted PANSS total scores were higher for fPIN-2 than fPIN-1 (t = 3.51, p < 0.001), mPIN-1 (t = 4.02, p < 0.001), and mPIN-2 (t = 2.55, p < 0.05).

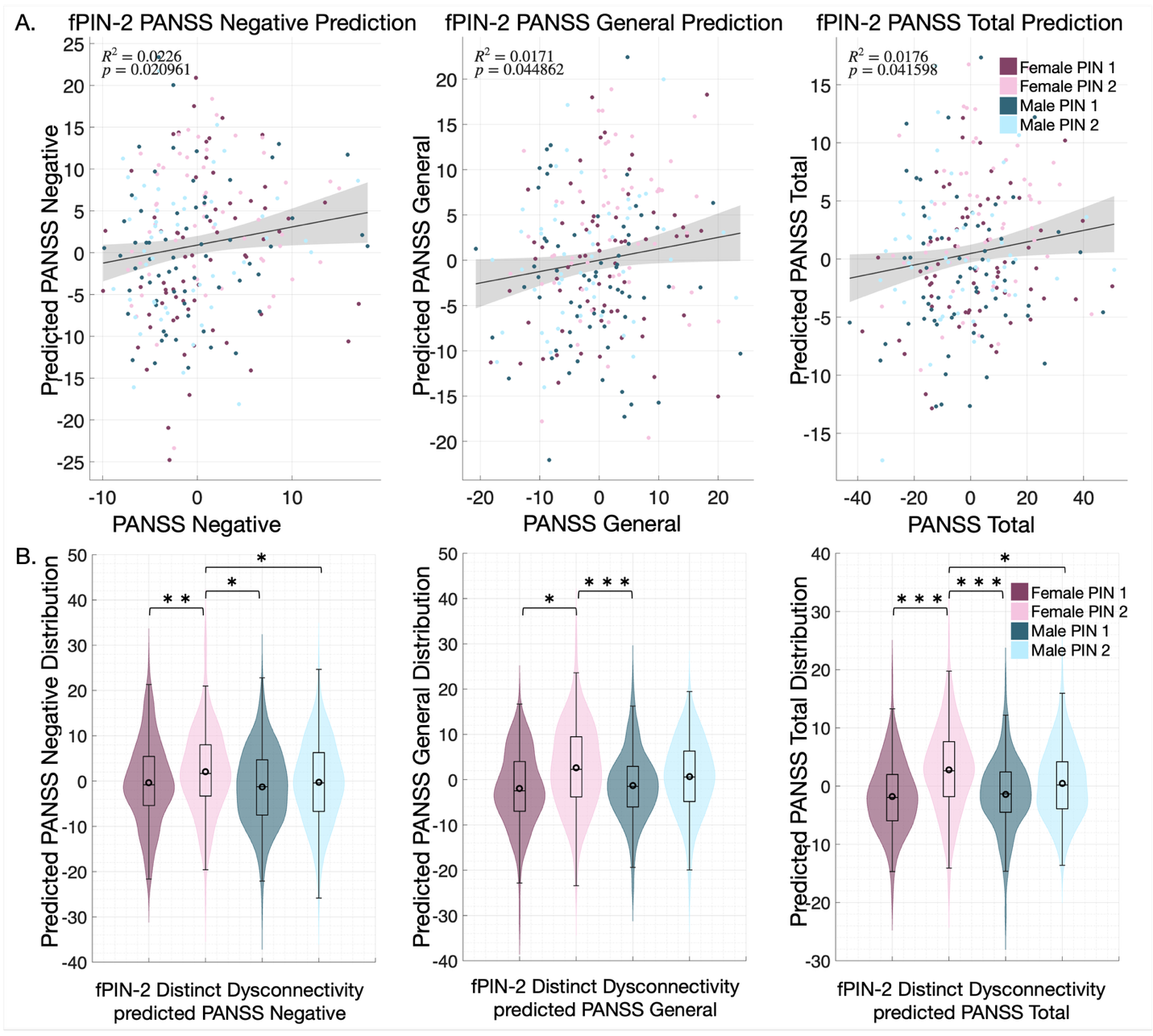

**Supplementary Figure 3: fPIN-2 disrupted subsystems predict PANSS symptom severity.** LASSO models trained using fPIN-2 disrupted subsystems were validated in the replication set. (A) Scatter plots show observed vs. predicted PANSS negative, general, and total scores with corresponding R² and p-values. (B) Violin plots compare predicted PANSS scores across the four PINs, demonstrating significant subtype differences (＊p < 0.05, ＊＊p < 0.01, ＊＊＊p < 0.001).

Supplementary Figure 4 illustrates the features contributing to the PIN-specific brain-BACS models and fPIN-2 PANSS models (Supplementary Figure 3). Together, these findings indicate that all PIN-specific disrupted subsystems are associated with distinct network pathways underlying cognitive impairment, while only the fPIN-2 subsystem showed a unique symptom pathway. These results suggest that clinical presentations may arise through subtype-specific neurobiological mechanisms.

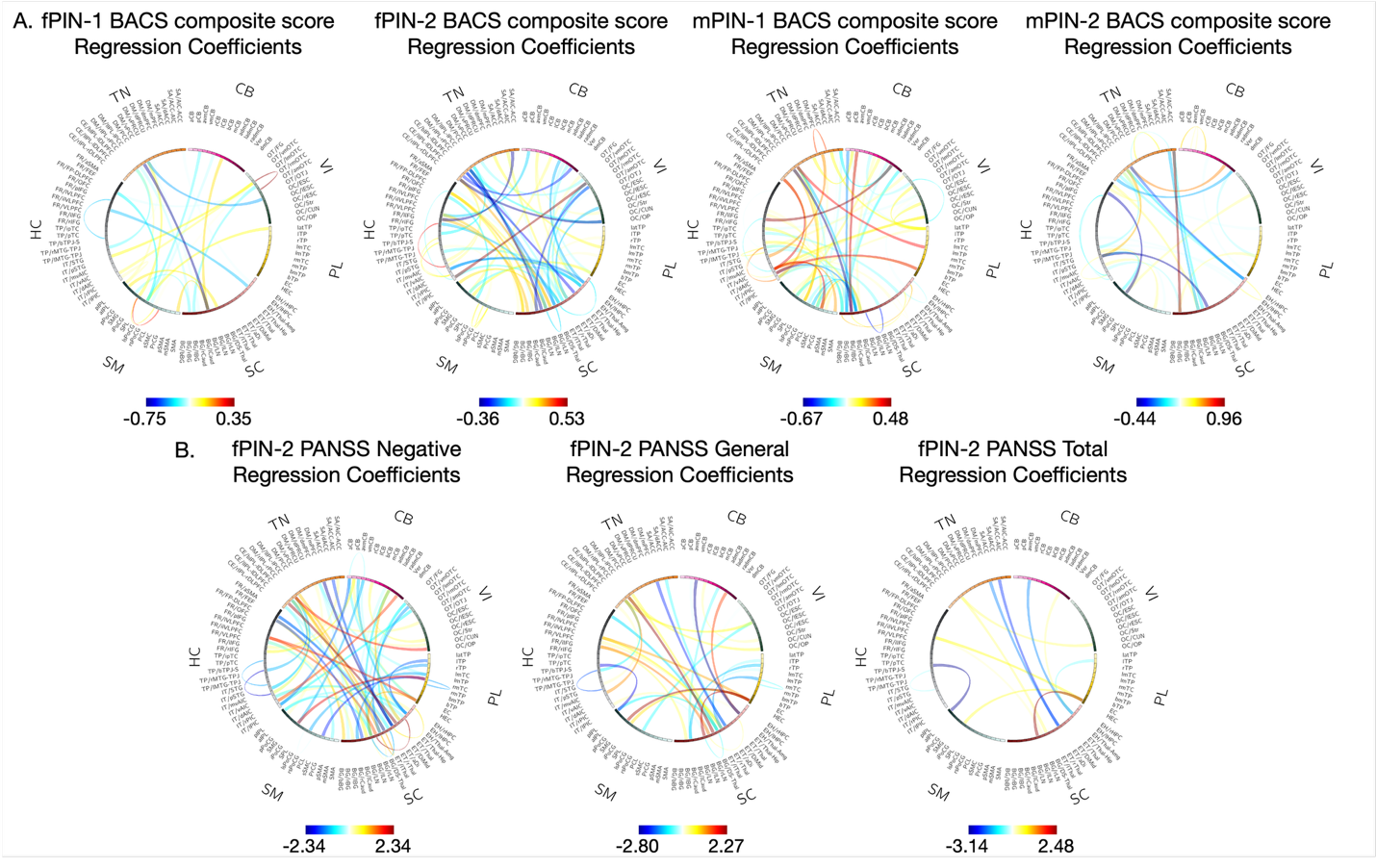

**Supplementary Figure.4: Sex-dominant Psychosis Imaging Neurosubtypes (PINs) underlying connectivity features predicting phenotype measures**. Connectograms display msFNC features selected by LASSO regression from PIN-specific disrupted subsystems, with links representing regression coefficients. A) Connectivity features contributing to Brain-Predicted Cognition models for fPIN-1, fPIN-2, mPIN-1, and mPIN-2. B) Connectivity features selected by the fPIN-2 subsystem for predicting PANSS negative, general, and total scores. Connectivity domains shown in connectograms include Cerebellar (CB); Visual (VI), comprising Occipitotemporal (OT) and Occipital (OC) subdomains; Paralimbic (PL); Subcortical (SC), comprising Extended Hippocampal (EH), Extended Thalamic (ET), and Basal Ganglia (BG) subdomains; Sensorimotor (SM); Higher Cognition (HC), comprising Insular Temporal (IT), Temporoparietal (TP), and Frontal (FR) subdomains; Triple Network (TN), comprising Central Executive (CE), Default Mode (DM), and Salience (SA) subdomains.
